# Sedimentary ancient DNA shows terrestrial plant richness continuously increased over the Holocene in northern Fennoscandia

**DOI:** 10.1101/2020.11.16.384065

**Authors:** Dilli P. Rijal, Peter D. Heintzman, Youri Lammers, Nigel G. Yoccoz, Kelsey E. Lorberau, Iva Pitelkova, Tomasz Goslar, Francisco J.A. Murguzur, J. Sakari Salonen, Karin F. Helmens, Jostein Bakke, Mary E. Edwards, Torbjørn Alm, Kari A. Bråthen, Antony G. Brown, Inger G. Alsos

## Abstract

The effects of climate change on species richness is debated but can be informed by the past. Here, we assess the impact of Holocene climate changes and nutrients on terrestrial plant richness across multiple sites from northern Fennoscandia using new sedimentary ancient DNA (*sed*aDNA) data quality control methods. We find that richness increased steeply during the rapidly warming Early Holocene. In contrast to findings from most pollen studies, we show that richness continued to increase through the Middle to Late Holocene even though temperature decreased, with the regional species pool only stabilizing during the last two millennia. Furthermore, overall increase in richness was greater in catchments with higher soil nutrient availability. We suggest that richness will rapidly increase with ongoing warming, especially at localities with high nutrient availability and even in the absence of increased human activity in the region, although delays of millennia may be expected.

## Introduction

Our ability to counter the current loss of biodiversity is dependent on how well we understand the causes of its change through time. However, the trajectory of biodiversity, especially in response to ongoing climate change, is vigorously debated (*1*, *2*), with discrepancy among short-term biodiversity patterns at global, regional, and local scales, whereby local processes may compensate or even counteract global trends (*3*). Our understanding of how species pools - the accumulated species richness at a given spatiotemporal scale (sensu *4*) - affect biodiversity patterns through time is limited, in part because constructing past species pools from present-day data is a non-trivial task (*5*).

The largest impact of ongoing climate change is expected to be at high latitudes (*6*). Field and modelling analyses suggest plant species richness will increase at high latitudes in Europe as summer temperature increases (*7*). Short-term observational studies, however, suggest that colonization by terrestrial species is lagging behind shifts in temperature isotherms (*8*), which can be compensated in the short term by local extinction lags (*1*). Therefore, studies addressing species pools and local richness at high latitudes and at different spatiotemporal scales are warranted to increase our understanding of biodiversity responses to ongoing climate change.

Changes in species richness by other drivers, such as nutrient levels, species introductions, and dispersal lags, are often context-dependent and hence difficult to predict. For example, edaphic variation, including variation in nutrient content, is hypothesized to strongly influence establishment, ecological drift, and niche selection, which all affect the local species pool, and this in turn affects richness (*9*). Experimental approaches have shown a non-linear impact of fertilization on Arctic plant richness and their ecosystem functions (*10*). An overall greater species richness has been reported from calcareous as compared with siliceous bedrock areas in the European Alps and northern Fennoscandia, whereby leaching of the former produces neutral to acidic microenvironments, providing a mosaic of habitats that may promote species establishment and an increase local richness (*11*, *12*). Human land use may also increase soil fertility and thereby richness (*13*), but the overall human impact at high latitudes in Europe is low (*14*). There is also evidence that the trajectory of succession, particularly soil formation, after glacier retreat varies due to abiotic rather than biotic factors (*15*). Furthermore, it has been found that regional plant species richness in previously glaciated regions may still be responding to past deglaciation, whereas local richness may be determined by local habitat factors (*16*).

There is a clear need for long-term data at the regional and local scales to better understand biodiversity trends (*17*). Paleoecological studies, especially those based on pollen (palynological) analyses, provide direct long-term evidence of plant biodiversity change and have been widely used to estimate effects of climate changes on richness (*18*–*20*). Contrasting richness patterns have been observed in different regions over the Holocene (11.7 thousand calibrated years before present (ka) to recent). In the European northern boreal region (Scotland, Fennoscandia, Iceland, Baltic States, NW Russia) pollen richness shows an overall decrease from 11.7-7.0 ka, followed by an increase to nearly peak levels by recent times (*19*). In the far north of Fennoscandia, however, a study spanning a forest to shrub-tundra gradient shows an inconsistent richness pattern through the Holocene (*21*). If plant dispersal had been slow, due to not being able to track a changing climate as it changes (dispersal lag) for example, then diversity in previously glaciated areas would be expected to increase over time. Such a scenario was not observed in central Sweden (*22*) but was inferred in Norway (*20*). These studies highlight the challenges of comparing pollen richness across different vegetation zones, which is confounded by inferences based on pollen being impacted by over-abundance of a few taxa (swamping), under-representation of other taxa, or low abundance of all taxa, the latter of which is a particular problem above the treeline (*23*).

An alternative, emerging proxy for examining long term, regional scale richness and species pool trends is sedimentary ancient DNA (*sed*aDNA). Compared to pollen, *sed*aDNA provides higher taxonomic resolution, has fewer problems with swamping, and is considered to only represent the local plant community (*24*–*26*). In a small catchment, *sed*aDNA may, therefore, also register the effect of drivers on a local rather than regional scale (*27*). Ground truthing studies show a strong correlation between modern sedimentary DNA-inferred richness and richness of modern vegetation around lake catchments (*28*). However, in contrast to pollen, *sed*aDNA studies have focused almost exclusively on single sites (but see *27*, *29*, *30*), thereby limiting our spatial understanding of richness and species pool patterns. A key challenge to combining multi-site *sed*aDNA data from lake cores is that data need to be directly comparable, both within (temporal) and between (spatial) sites, otherwise biased richness estimates resulting from data quality problems could potentially lead to erroneous local- and regional-scale inferences.

Here, we generate the largest *sed*aDNA data set to date, consisting of 387 dated samples from 10 sites in northern Fennoscandia using vascular plant metabarcoding, and harmonize the entire data set using standardized taxonomy and novel data quality control measures that will be highly applicable to the wider environmental DNA community. Capitalizing on this harmonized high resolution taxonomic data set, we estimate richness, local species pools (accumulated richness per catchment), and the regional species pool (accumulated richness for all 10 sites) throughout the Holocene which we compare to two potential drivers, climate and local edaphic conditions. We find that temperature and soil nutrients are important drivers, but suggest that dispersal lags and habitat diversification also provide an important mechanism for plant richness changes through time. By providing refined paleoecological insights, sedaDNA data are well positioned to increase the precision of integrative ecological models for predicting the consequences of ongoing climate change.

## Results

### Age-depth models

We constructed Bayesian age-depth models for 10 lake sites (Figs. 1, 2, S1) to estimate the age of each individual *sed*aDNA sample. Since the cores were all central or near-central lake locations and the lakes were medium to small with in most cases only one depositional basin, the age-depth curves were approximately linear or curvilinear with three exceptions (Figs. 2, S1). Kuutsjärvi had a distinct reduction in sedimentation rate from around 4.0 ka. Sandfjorddalen had stepped changes in the sedimentation rate with possible hiatuses in the Early Holocene (11.0-8.0 ka) and Late Holocene between 6.0 and 2.0 ka. This probably reflects its position in the valley floor as a flow-through lake. Lastly, Sierravannet had a distinct upturn in the accumulation rate around 0.6 ka to present, which occurs after a putative flood event at ca. 48-40 cm composite depth (equivalent to ca. 0.6 ka). This 8 - cm layer is characterised by a dark band in the visible stratigraphy, a rapid decrease and then increase in organic carbon (loss-on-ignition, LOI), and two older-than-expected dates, which were consequently not included in the age-depth model. Given that this lake has the largest catchment area and there is a change in lithology, this is probably the result of flooding from the upstream lakes and fluvial network. The terrestrial plant taxonomic richness trends are unaffected by the removal of the four *sed*aDNA samples that fall within this flood event window (Fig. S2). For the interpretation of the *sed*aDNA records, the age-depth models provided similar temporal resolution of 158-616 years per sample for all lakes except Sierravannet, which had 63 years per sample. Six of the sedimentary records covered the entire Holocene (Figs. 2, S1) and all except one (Sierravannet) covered the three periods of the Holocene (Early, 11.7-8.3 ka; Middle, 8.3-4.25 ka; Late Holocene, 4.25-0.0 ka), although the usable Nesservatnet record was reduced to the Late Holocene after removal of low quality *sed*aDNA samples (see below). For two records that extend into the lateglacial (late Pleistocene; Langfjordvannet, Nordvivatnet), the age-depth models are not well constrained in the lateglacial period. For Langfjordvannet, three hiatuses were inferred by Otterå (*31*). We tentatively infer a single hiatus for Nordvivatnet (Fig. 2), which could have been caused by a rock slide, and is based on the recovery of glacially-scarred stones (impressions shown in visible stratigraphy in Fig. S1H), low organic carbon (LOI), and comparable radiocarbon dates.

**Fig. 1.**
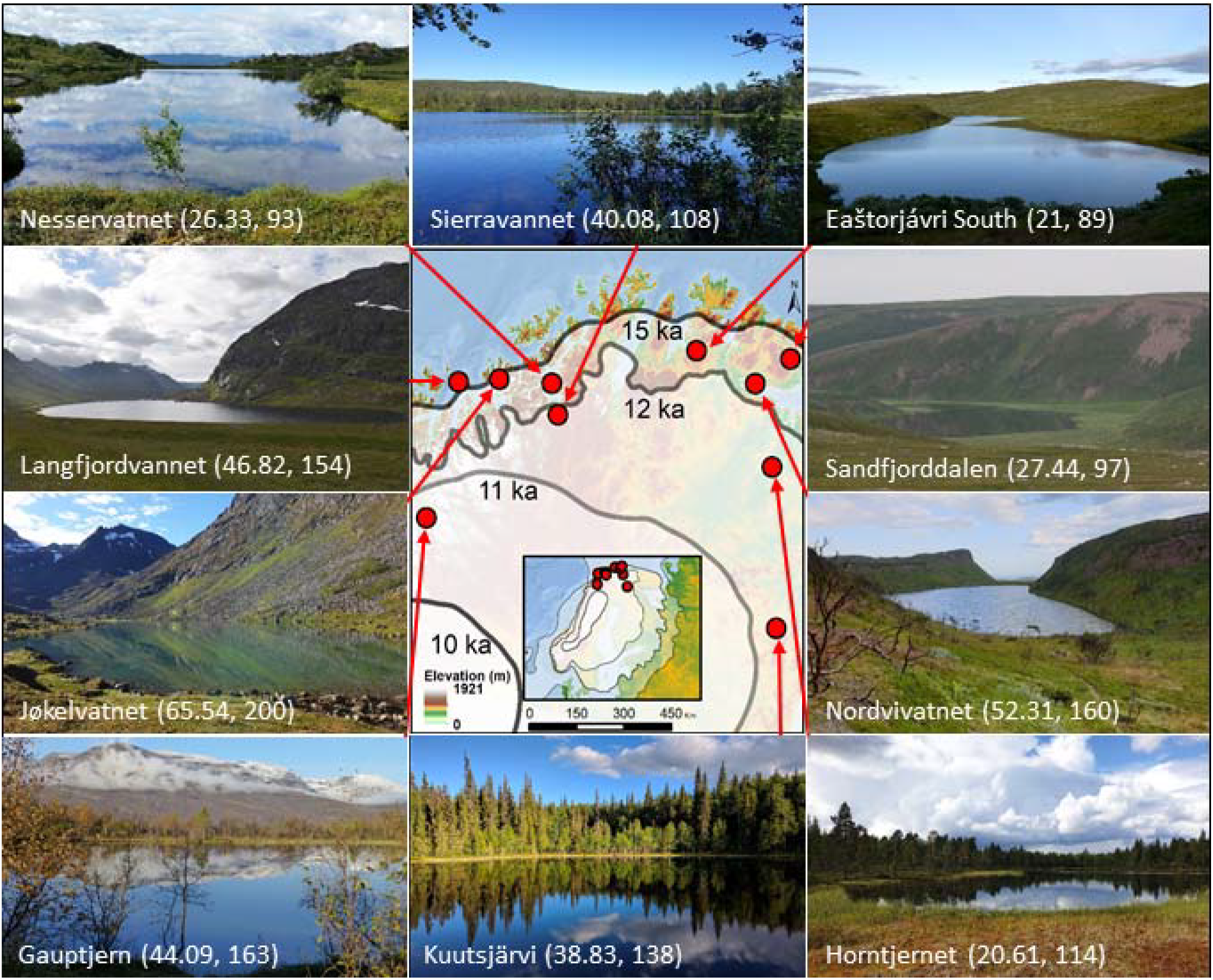
Geographical setting of the study area and images of the ten lakes. The extent of the Scandinavian ice sheet (the most credible extent of *79*) at 21.0 (inset only), 15.0, 12.0, 11.0, and 10.0 ka are indicated by semi-transparent layers. Lake names are followed by mean taxonomic richness and total taxa recorded in each lake (local species pool). See Data set S1 for further site information. Map data source: European Environment Agency; photo credits: Jøkelvatnet, Lasse Topstad; Sandfjorddalen, Leif Einar Støvern; Langfjordvannet & Eaštorjávri South, Dilli P. Rijal; Kuutsjärvi, Karin Helmens; all others, Inger G. Alsos.

**Fig. 2.**
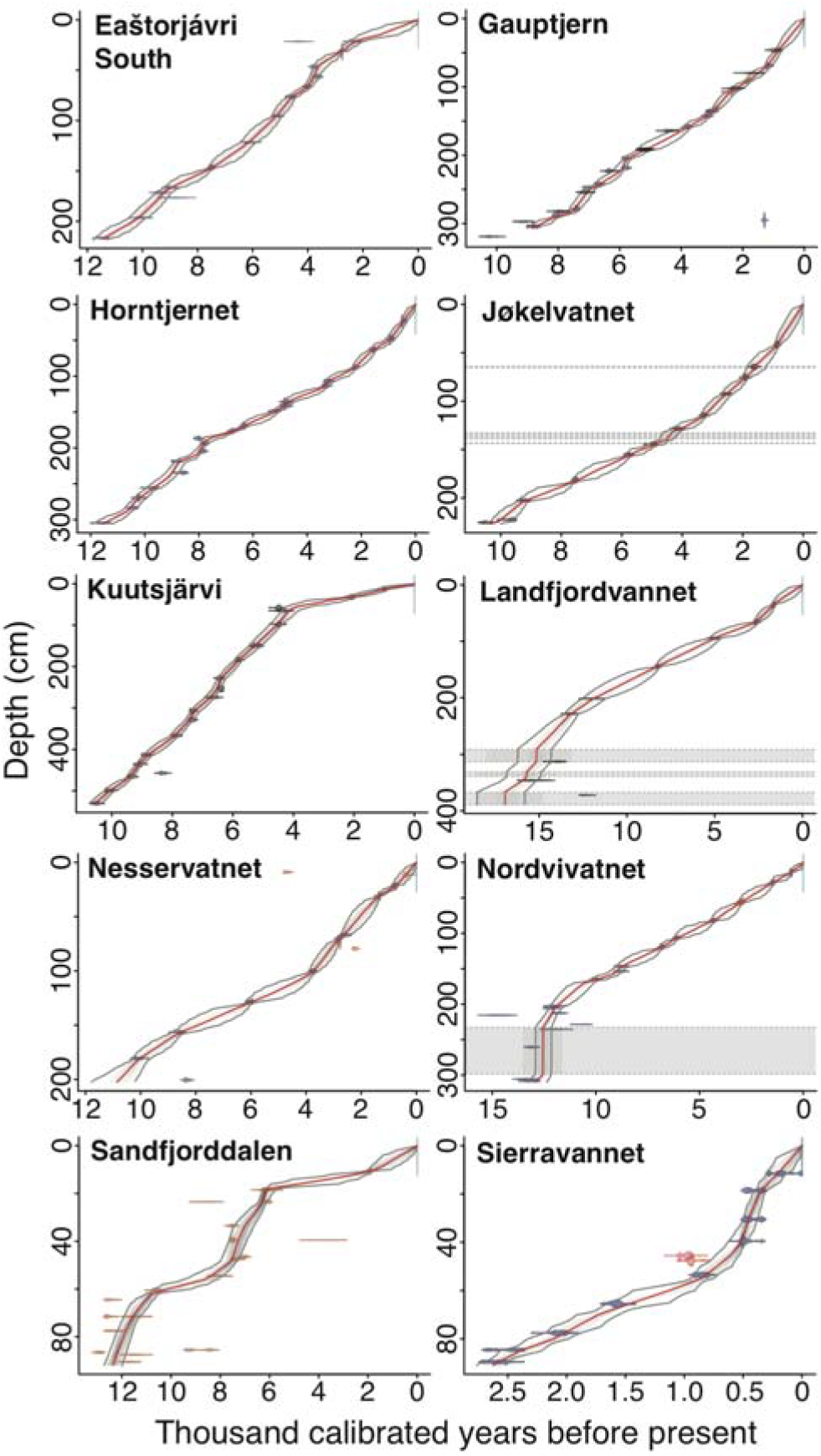
Bayesian age-depth models for ten lakes from northern Fennoscandia. Colors for calibrated radiocarbon dates follow Fig. S1. Excluded dates are in red, and inferred slumps are shown with grey shading. Median modelled ages are indicated by the redlines, with the bubbles representing 95% credibility intervals.

### Sedimentary ancient DNA data quality assessment

Across our 10 lake sediment records, we generated 91.6 million raw sequence reads from 387 sediment samples and 90 control samples. We observed notable differences in data quality between samples both within and between lake records during preliminary data exploration. We therefore sought to develop criteria to scrutinize the quality of samples and provide a cut-off for removing those considered problematic and which may have been impacted by poor DNA quality or methodological issues, such as extract inhibition. We developed two statistics based on the most read-abundant sequences, with the rationale that if these sequences were not replicable, then we cannot exclude methodological problems. We first developed a *metabarcoding technical quality (MTQ)* score to assess metabarcoding success on a per-sample basis. This score is the proportion of positive PCR detections across the 10 most read-abundant sequences within a sample prior to any taxonomic identification. We next developed a *metabarcoding analytical quality (MAQ)* score to assess the success of recovering sequences of interest. This score is the same as the *MTQ* score, except that the 10 most read-abundant and taxonomically-identified sequences, after removing those that matched the *blacklists*, were used. Sources of *MTQ* and *MAQ* score divergence include the co-amplification of unidentified and/or contaminant sequences. For all samples across the data set, we examined the distribution of *MTQ* and *MAQ* scores and used these to infer that samples should require an *MTQ* score of ≥0.75 and *MAQ* score of ≥0.2 to pass quality control (QC) and be included in downstream analyses (Figs. 3, S3, S4). This cutoff excluded all negative controls, which had an *MTQ* score of <0.75 and *MAQ* score of <0.1.

**Fig. 3.**
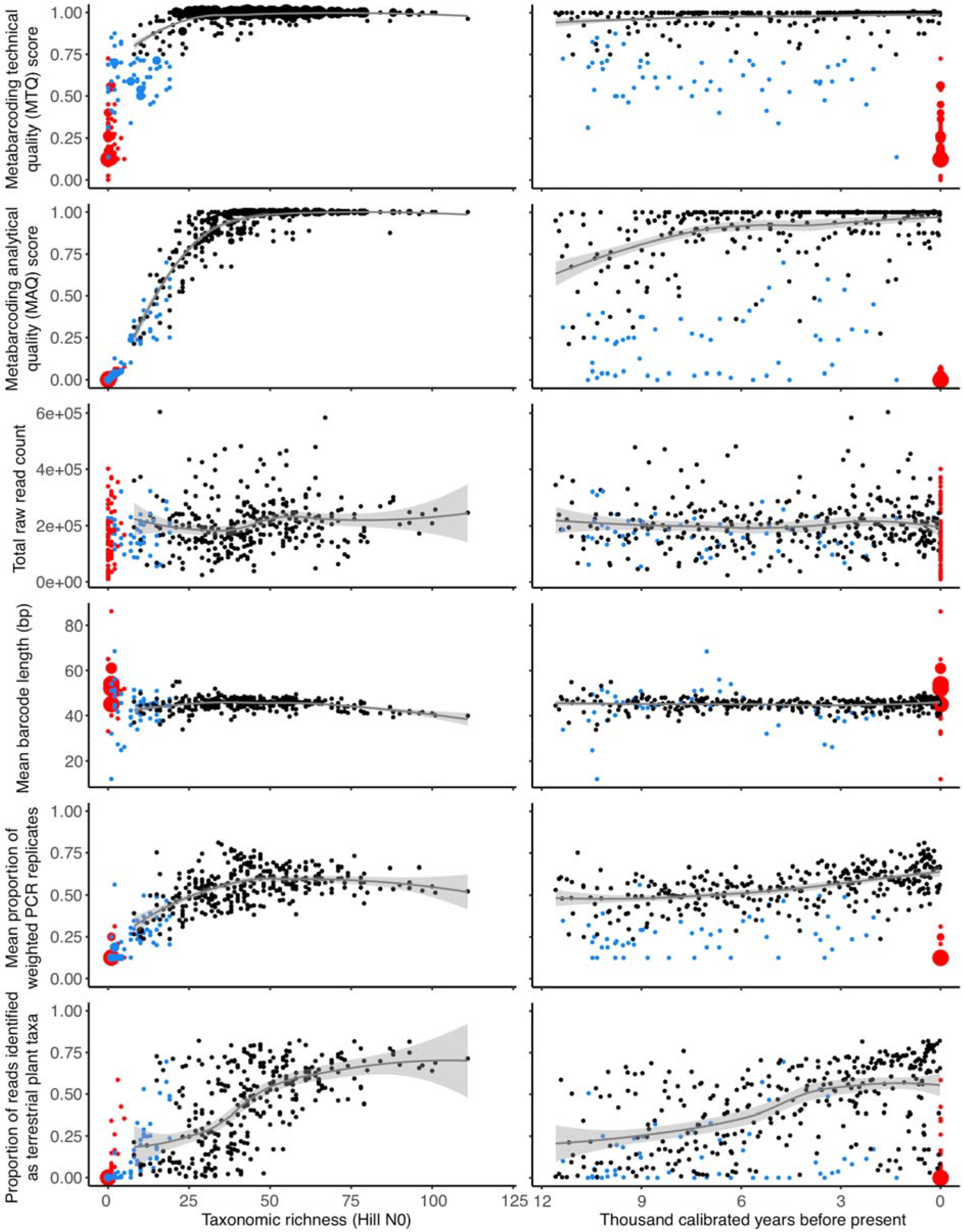
Correlations between observed taxonomic richness (left panels) and time (right panels) against six measures of *sed*aDNA data quality. Sample age minimally impacts *metabarcoding technical quality (MTQ)* and *metabarcoding analytical quality (MAQ)* scores, total raw read count, mean barcode length, and the mean proportion of weighted PCR replicates (*wtRep*) across the entire data set, although the proportion of reads identified as terrestrial plant increases through time. Data in black, samples that passed quality control (QC); blue, samples that failed QC; red, negative controls. Fitted loess-smoothed lines along with one standard error envelope are for samples that passed QC. Data for individual lakes are presented in Fig. S4.

After applying our QC thresholds and removing duplicates, we retained 316 samples (Fig. 3). This resulted in 12-55 samples retained per record (Data sets S4, S8; Fig. S3). We retained 402 barcodes, which were collapsed to 346 taxa with between 89-200 taxa recorded from each lake record (Table S1). Of these, 50% could be assigned to the species level (Data sets S6, S7). As our focus was on the terrestrial plant diversity, we excluded 13 algae and 36 aquatic plant taxa. Nine taxa were only present in samples that failed QC. Thus, our final dataset retained 288 terrestrial plant taxa detected in 316 samples.

We next explored the relationship between observed taxonomic richness and/or sample age against six measures of *sed*aDNA quality, with each measure calculated on a per sample basis.

#### MTQ and MAQ scores

Both scores correlate with richness when richness is low (<25-30; Fig. 3), which is likely an artifact of the requirement that the 10 best represented barcodes are required to calculate these scores. At higher richness values, both *MTQ* and *MAQ* scores are uniformly high. *MTQ* scores are minimally impacted by sample age (Figs. 3, S4A), although samples older than ~8.0 ka tend to have lowered *MAQ* scores (Fig. 3), which is driven by the Eaštorjávri South, Langfjordvannet, Kuutsjärvi, Nesservatnet, and, to a lesser extent, Jøkelvatnet records (Fig. S4B).

#### Raw read count summed across PCR replicates

Richness may be influenced by differences in total sequencing depth between samples, whereby we would expect increased total depth to correlate with richness as the likelihood of detecting read-rare taxa is increased (e.g. (*32*)). However, we do not observe a relationship between richness and raw read count (Fig. 3), suggesting that differences in sequencing depth do not influence richness in our data set. This is consistent with the results of the rarefied richness analyses (Table S2). We also do not observe a relationship between sample age and sequencing depth (Figs. 3, S4C), suggesting there is no temporal or stratigraphic bias in our ability to generate raw reads. We note that samples that passed or failed QC had comparable total sequencing depths.

#### Mean length of identified barcodes through time

As ancient DNA fragments into shorter molecules over time (*33*), a reduction in mean barcode length with sample age may suggest that longer barcodes are no longer preserved thus biasing estimates of temporal richness patterns. However, this assumes that barcode length is independent of taxonomic group and/or ecological community composition, which may not always be the case. We do not observe a decrease in mean barcode length with age in samples that pass QC (Figs. 3, S4D). Samples that fail QC are often mean length outliers that show no relationship to age, but are rather an artifact of small sample sizes. Minor decreases in mean barcode length with sample age are observed for Kuutsjärvi and Langfjordvannet (Fig. S4D).

#### Mean proportion of weighted PCR replicates (wtRep)

The mean *wtRep* provides a measure of mean taxon detectability in samples. If barcode template concentrations in *sed*aDNA extracts are low, then we would expect recovered richness to be limited, due to dropout, with a corresponding reduction in detectability of taxa that are observed. Consistent with this expectation, we find that mean *wtRep* values are comparable for samples with richness >25-30 (Fig. 3), but that mean *wtRep* and richness are correlated when richness is <25-30. However, we observe only a modest decrease in mean *wtRep* values with sample age (Fig. 3), suggesting that taxon detectability is not affected by age. The greatest age-related reductions in mean *wtRep* values are in samples from Langfjordvannet, Kuutsjärvi, Nesservatnet, and, to a lesser extent, Jøkelvatnet (Fig. S4E). We note, *a posteriori*, that mean *wtRep* and *MAQ* scores may not be independent measures, especially for samples with low richness (<25-30).

#### The proportion of raw reads assigned to terrestrial plant taxa

If there is co-amplification of non-terrestrial plant, algae, contaminant, and/or other off-target molecules, then terrestrial plant richness may be reduced by swamping. Interestingly, we observe that at least ~40% of reads are required to be identified as terrestrial plant taxa for observed richness to exceed ~60 taxa (Fig. 3). When richness is <60, we observe that the proportion of reads identified as terrestrial plant taxa decreases with richness (Fig. 3). However, we note that there is large variance around this trend and so, for example, samples from the Middle Holocene of Sandfjorddalen have comparable observed richness to Early Holocene samples from Gauptjern (Fig. S3), even though reads from the former are swamped by reads from aquatic plant taxa (Fig. S5). Across the entire data set, there is a gradual reduction in the proportion of reads identified as terrestrial plant taxa with sample age, which is driven by the Eaštorjávri South, Kuutsjärvi, and Nesservatnet records (Figs. 3, S4F, S5). Samples that failed QC tended to have a lower proportion of reads identified as terrestrial plant taxa (Figs. 3, S4F, S5).

Overall, we found that the *MTQ* and *MAQ* score QC thresholds removed the worst performing samples. The records with the best *sed*aDNA quality are Gauptjern, Horntjernet, Nordvivatnet, Sandfjorddalen, and Sierravannet. Samples from the Early Holocene should be treated with more caution from the Eaštorjávri South, Kuutsjärvi, Langfjordvannet, and Jøkelvatnet records (Figs. S4, S5).

### Local richness and species pool

To evaluate how richness and the local species pool changed through time, we calculated the accumulated and observed number of taxa in each sample within each lake as measures of local species pool and richness respectively. The local species pools increased over time for all catchments (Fig. 4) with the highest recorded at Jøkelvatnet (200 taxa), which today drains a catchment that has a Late Holocene glacier in its upper reaches (Fig. 1; Supplementary Materials, SI1). Rich species pools were also found at Gauptjern, which is at the border between pine and birch forest, and at Nordvivatnet and Langfjordvannet, which have a mixture of heathland, birch forest, and scree slope in their catchments. Somewhat lower species pools were found at the two sites in pine forest, Horntjernet and Kuutsjärvi, and at Sierravannet, a site with birch forest, and pine and larch plantations. The two shrub-tundra sites, Eaštorjávri South and Sandfjorddalen, had smaller species pools, similar to Nesservatnet, which is surrounded by heathland/mires (93 taxa) and located on the small island of Årøya.

**Fig. 4.**
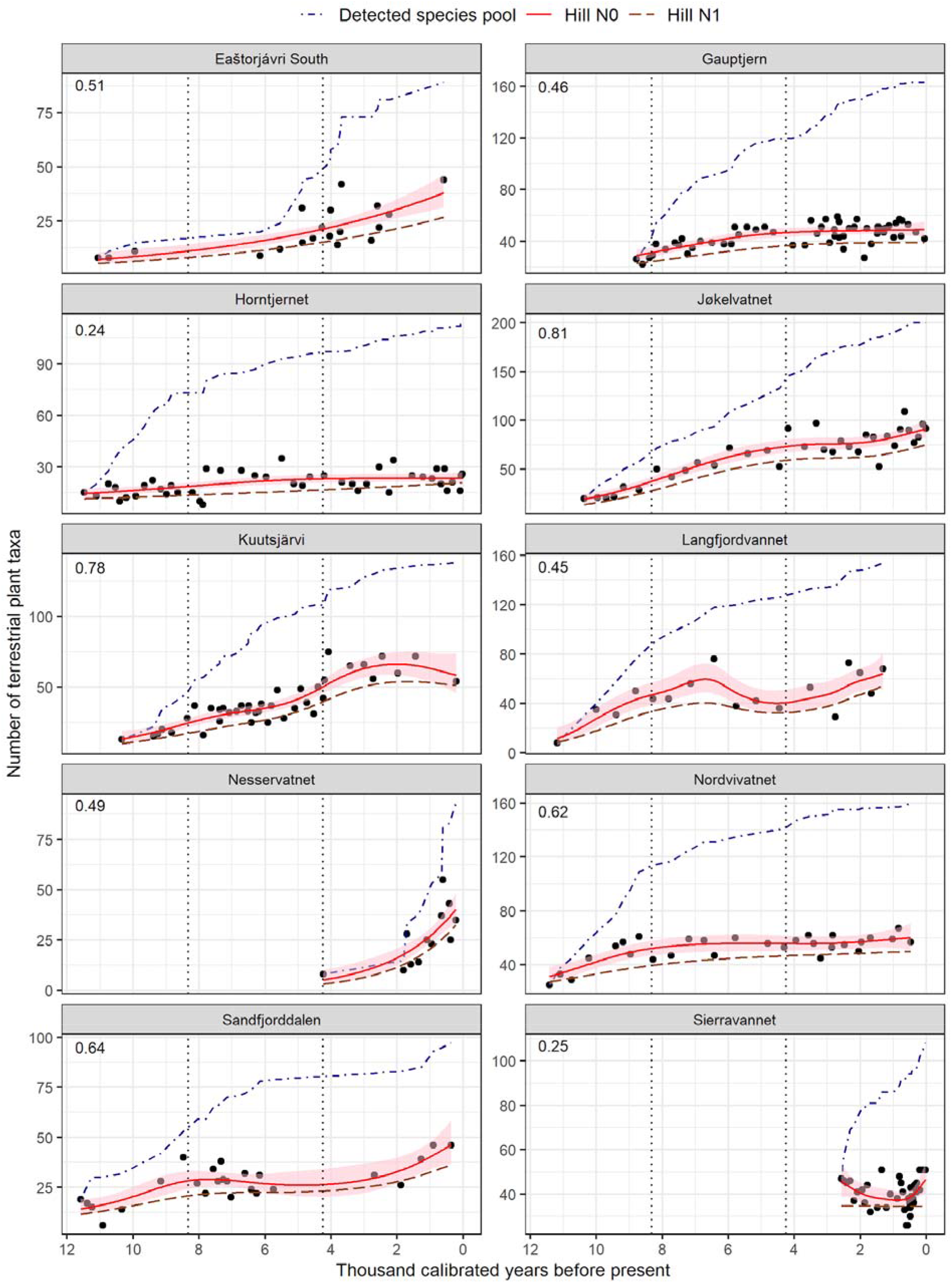
Holocene patterns of local terrestrial plant richness in 10 lake catchments from northern Fennoscandia. The predicted values for observed taxonomic richness (Hill N0) are based on the generalized additive mixed models (GAMM; solid red line) along with 95% confidence intervals in pink shading. Numbers in the top-left corner of each plot represent adjusted R-square values for the GAMMs. The fitted lines for Hill N1 are indicated by a dashed brown line. The development of local species pools are expressed in terms of detected cumulative count of taxa (blue dot-dashed line) through time. The Early (11.7-8.3 ka), Middle (8.3-4.25 ka), and Late Holocene (4.25-0.0 ka) periods are indicated by vertical dotted lines. Note difference in scale on the y-axes.

There were clear differences among lakes, both in the overall levels of richness and in the change in richness over the period (Fig. 4). The mean (±SD) taxonomic richness (Hill N0) ranged from 20.6 (± 6.4) at Horntjernet to 65.5 (± 24.5) at Jøkelvatnet, whereas Hill N1 ranged from 14.9 (± 7.8) at Eaštorjávri South to 52.4 (± 20.5) at Jøkelvatnet (Fig. 4; Table S1). The rarefied richness based on read count showed a strong correlation with observed taxonomic richness (R=0.82-0.99; Table S2), suggesting that the observed pattern was not affected by sequencing depth. The Hill N1 (common taxa) showed temporal patterns that mirrored those of observed taxonomic richness for all the lakes except Sierravannet (Fig. 4).

We observed a significant effect of the age of samples on taxonomic richness as indicated by statistically significant smooth terms in generalized additive mixed models (Table S3), except for Sierravannet, which only covered 2.6 ka, and where diversity suddenly dropped around 0.6 ka, corresponding to a putative flood event (Fig. S2). For two of the lakes, Eaštorjávri South and Nesservatnet, a near linear pattern of increase in taxonomic richness through time (edf=1) was recovered. On the other hand, Langfjordvannet had the most complex pattern of increase in richness (edf=5.93, Table S3). The steepest increase was seen in the Early and Middle Holocene for most lakes. Only at three sites, Nordvivatnet, Horntjernet and Gauptjern, did richness reach a plateau during the Late Holocene; for most lakes no levelling off was observed suggesting that richness is still increasing (Fig. 4).

### Regional richness and species pool

To assess temporal patterns of regional species pool and richness, we calculated the accumulated number of detected taxa from the entire quality controlled data over the Holocene as a measure of the regional species pool, and the number of taxa detected in each sample as a measure of richness (Figs. 5A, S6). During the Early Holocene, there was a strong increase in the regional species pool size. The regional species pool increased monotonically from the Early Holocene to ca. 7.0 ka, the rate slowed in the period ca. 7.0-5.0 ka followed by an uptick from 5.0-3.3 ka before stabilising. The regional species pool levelled off over the last two millennia with an increase of just 10 taxa (3.5% of the total).

**Fig. 5.**
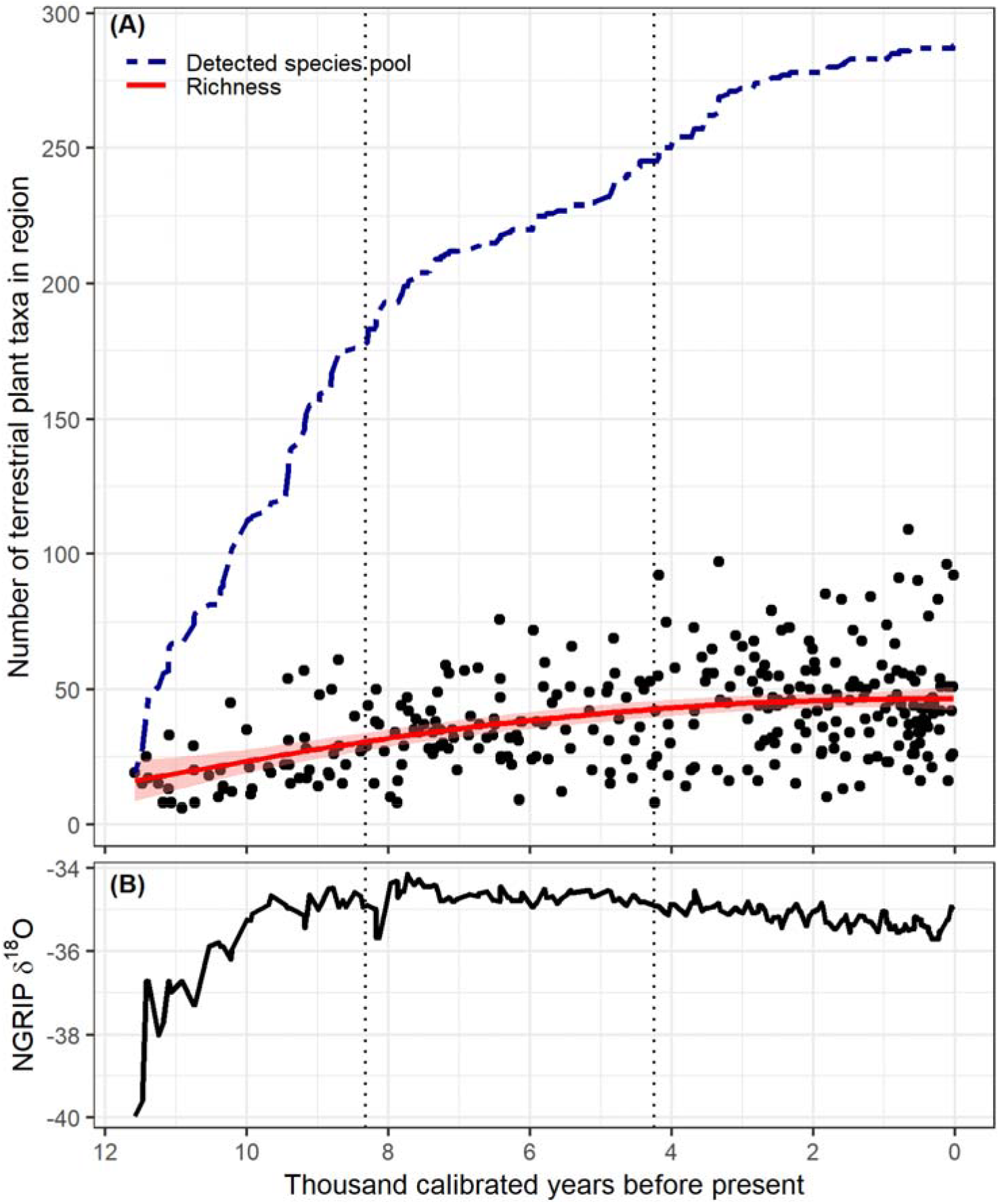
The accumulated regional species pool, taxonomic richness of each sample, and NGRIP temperature proxy across the Holocene of northern Fennoscandia. (**A**) accumulation of the detected regional species pool (defined as cumulative number of taxa; double-dashed line) as well as number of taxa detected per sample (n=316) along with the 95% confidence interval (pink shading) of the fitted line (solid red line) based on a generalized additive model, and (**B**) variation in temperature reflected by North Greenland Ice Core Project (NGRIP) δ^18^O values (*80*), with temperature being positively correlated with δ^18^O values. The overall patterns remain the same if two shorter cores spanning only the Late Holocene are excluded (Fig. S6). The Early (12.0-8.3 ka), Middle (8.3-4.25 ka), and Late Holocene (4.25-0.0 ka) periods are indicated by dotted vertical lines.

The mean (±SE) predicted taxonomic richness (Hill N0) based on a generalized additive model showed a steep increase during the Early Holocene from 13.8±3.9 to 31.8±1.5 taxa per sample when evaluated using 500-year time windows. Richness continued to increase during the Middle Holocene (33.4±1.5 to 42.7±1.3), and showed only a minor increase during the Late Holocene (43.7±1.3 to 45.9±2.0).

### Richness in relation to local and regional species pools

To assess how local and regional species pool affects richness (beta-diversity) at respective scales, we used the accumulated number of taxa within a lake and 500-year time bin as measures of local and regional species pools respectively, and mean number of taxa at respective scales as richness estimates. There was a strong positive association between mean terrestrial plant richness and the local species pool of lakes, where 82% of the variation in local richness was explained by local species pool (R^2^_adj_ =0.82, p<0.001, df=8; Fig. 6A). The mean richness of lakes represented about 24% to 37% taxa of the local species pool, except at Horntjernet, where richness represented only 18% taxa of the local species pool. The mean local richness increased by nearly four taxa (regression slope=0.36) for every 10 taxa that were added in the local species pool. Similarly, we found a strong positive correlation between mean richness per 500-year time period and total taxa available in the respective period, and 86% of the variation in richness was explained by the regional species pool (R^2^_adj_ =0.86, p<0.001, df=21; Fig. 6B) where ~23% to 39% of the taxa from the regional species pool were represented by the mean richness. The mean regional richness increased by more than two taxa (regression slope=0.23) with the addition of 10 taxa in the regional species pool.

**Fig. 6.**
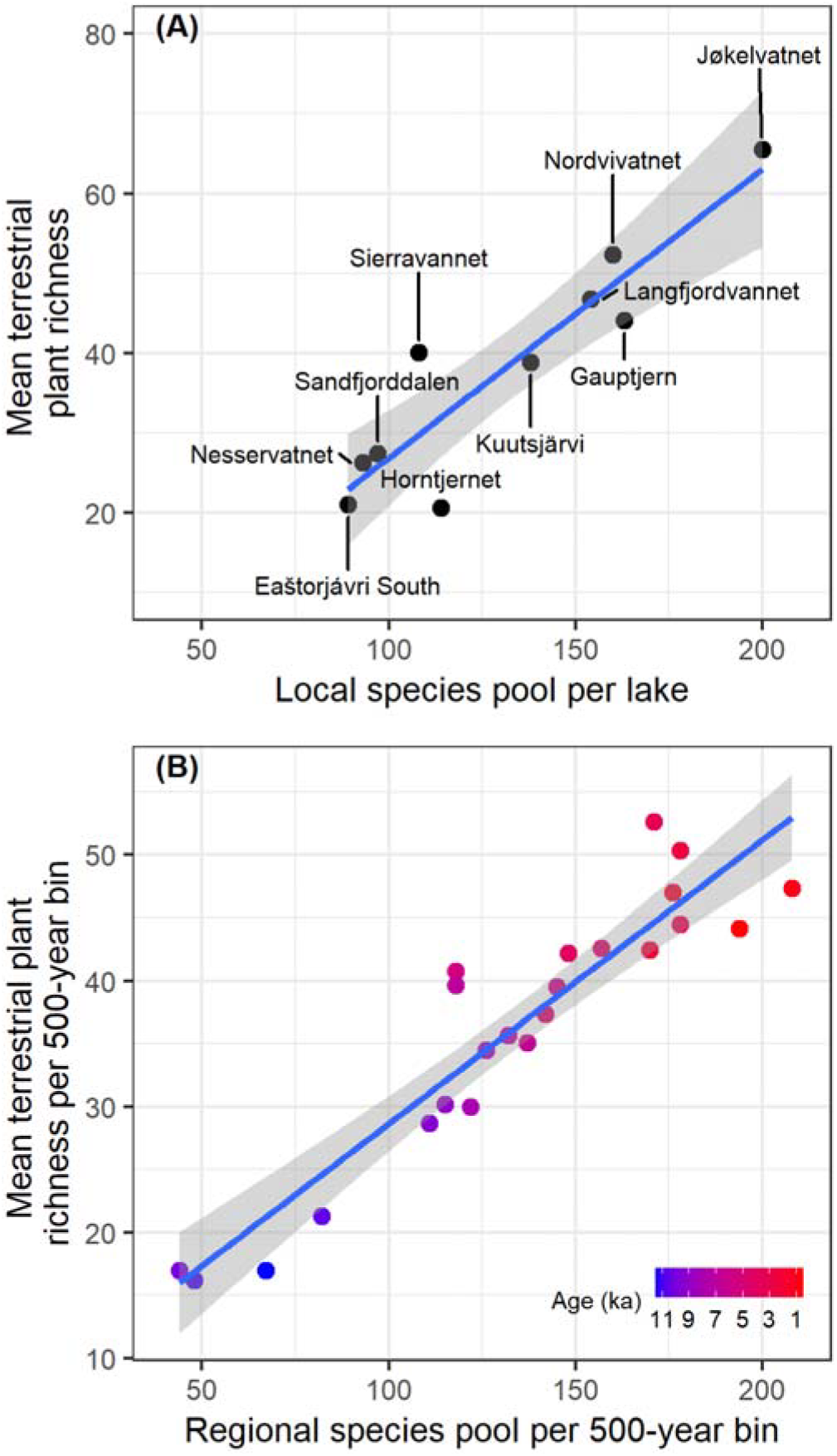
Relationship between species pool and taxonomic richness. Correlation between (**A**) local species pool (total number of taxa observed over the Holocene for individual lakes) or (**B**) regional species pool (total number of taxa observed per 500-year time bin) and the respective mean taxonomic richness of terrestrial plants in northern Fennoscandia.

### Impact of regional climate on plant richness

We used oxygen isotope (δ^18^O) values from the North Greenland Ice Core Project (NGRIP) (Andersen *et al.*, 2004) as a proxy for temperature to assess the impact of regional climate on local richness during different periods of the Holocene. Climate had a significantly positive effect on richness in the Early Holocene (β=0.23, SE=0.05, p<0.001), a marginal negative effect in the Middle Holocene (β=−0.26, SE=0.13, p=0.048), and a clear negative effect in the Late Holocene (β=−0.53, SE=0.12, p<0.001; Fig. 7A). In the Early and Late Holocene, temperature changed linearly through time at a rate (±SE) of 0.92 (±0.07) and −0.13 (±0.01) δ^18^O/ka respectively, whereas there was no directional change in temperature across the Middle Holocene (Fig. 5B).

**Fig. 7.**
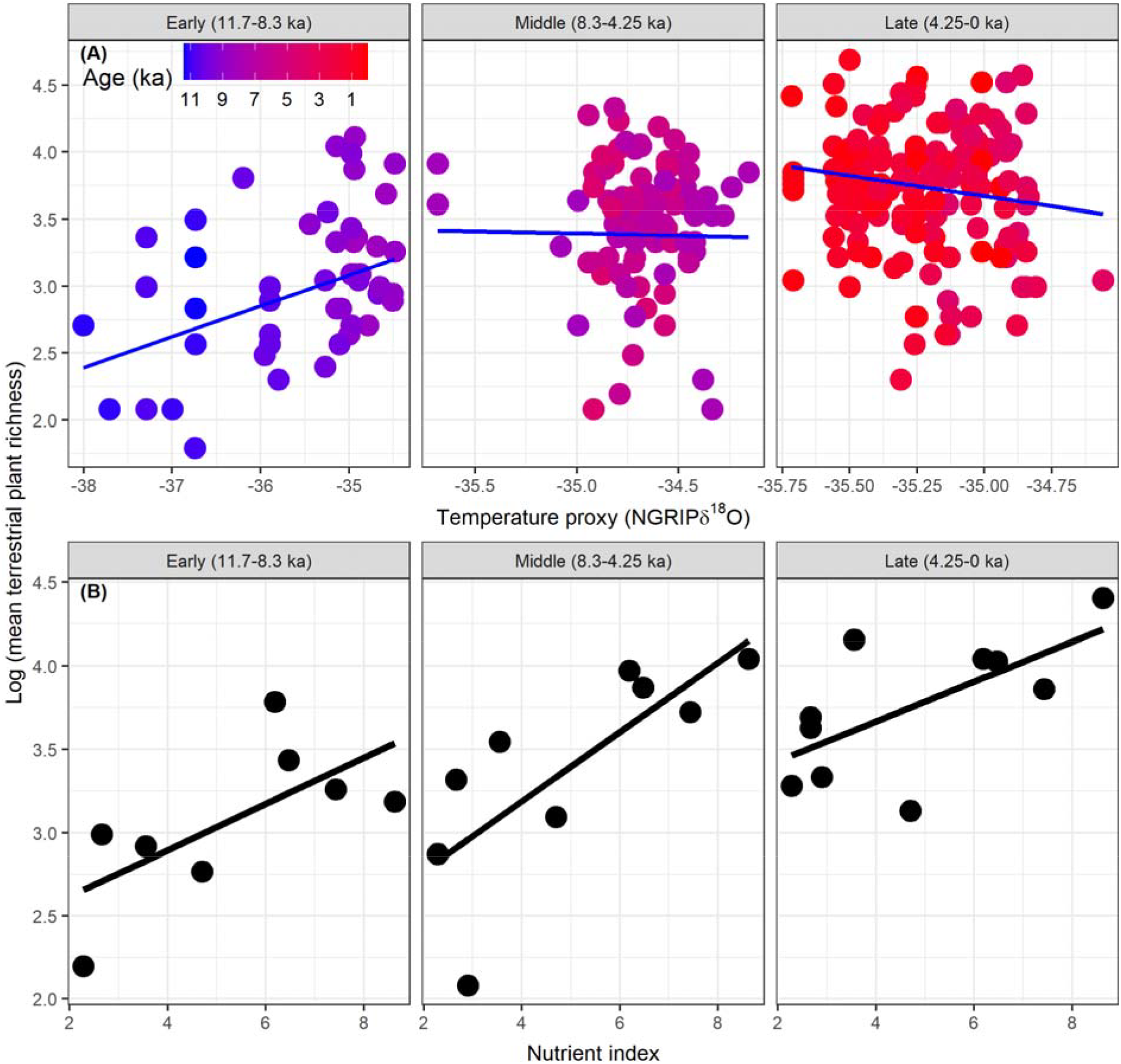
Impact of climate and nutrient index on observed terrestrial plant taxonomic richness. (**A**) Linear mixed effect model showing the impact of regional climate on taxonomic richness of terrestrial plants for three periods of the Holocene. Temperature is positively correlated with δ^18^O values. Two samples with NGRIP δ^18^O lower than −39 were not included in the analysis. Note difference in scale on x-axes. (**B**) Linear models showing spatial patterns of mean taxonomic richness of terrestrial plants with the nutrient index for three periods of the Holocene.

### Effect of nutrient availability on plant richness

We used a new semi-quantitative nutrient index based on rock type and its weatherability to assess how nutrient availability in the catchment area affects richness. We observed a positive correlation between nutrient index and taxonomic richness for all three time periods (Fig. 7B) although this was not significant for the Early Holocene (R^2^_adj_ =0.36, F(1,6)=4.92, p=0.07), which also had the smallest sample size. The nutrient index explained 51% and 35% of the variation in richness for the Middle (R^2^_adj_ =0.51, F(1,7)=9.22, p=0.02) and Late Holocene (R^2^_adj_ =0.35, F(1,8)=5.92, p=0.04) respectively. The nutrient-richness relationship was stronger for the Middle Holocene (β=0.21, SE=0.07, p=0.02) than the Late Holocene (β=0.12, SE=0.05, p=0.04). The effect of nutrient index on taxonomic richness was strongest when the impact of climate was negligible during the Middle Holocene. This suggests that a significant cause of site-to-site variation and sub-regional richness patterns was soil nutrient availability, which is dependent upon the bedrock and rate of weathering.

## Discussion

### The ability of *sed*aDNA to capture plant taxonomic richness

The mean observed taxonomic richness of terrestrial plants per sample and site (~21-66) is higher than that recovered for northern boreal sites based on pollen analyses (~20 taxa, *19*), but similar to pollen estimates from the Alps and Mediterranean (~30 taxa, *19*). The detected richness values are within the range that has been found in other recent studies of *sed*aDNA from northern sites (20-70 taxa per sample; *27*, *34*), although some shrub tundra (~13 per sample, *35*) and High Arctic (5-30 per sample, *36*) sites have notably lower estimates. Nevertheless, our results are consistent with other *sed*aDNA analyses that detect more taxa than pollen counts (*24*, *25*, *27*). Together with improved geographic fidelity due to a local signal, *sed*aDNA thereby improves our understanding of the spatial patterns and scale dependency of past plant diversity.

The temporal patterns evaluated here rely on the assumption that our ability to detect plant taxa in *sed*aDNA is not impacted by differential preservation, due to sample age or methodological problems such as DNA extract inhibition. Here we discarded samples of poor quality, that had metrics comparable to negative controls and thus may have been affected by methodological problems, and we broadly examined the quality of the retained samples. Half of our sites showed no evidence of declining *sed*aDNA quality with sample age, whereas the remainder had reduced quality in the Early Holocene interval. That our samples generally exhibited good *sed*aDNA quality throughout the study interval is likely due to a combination of excellent DNA preservation in the cold environments of high latitudes (*37*) and the young age of the samples (<11.7 ka) relative to the known upper limit of aDNA preservation (~560-780 ka, *38*). As multi-site *sed*aDNA studies become common, it will be crucial that data quality is scrutinized and, where possible, standardized to allow for biologically meaningful comparisons between sites.

### Nutrient availability and plant richness

In considering the positive association between nutrient index and mean taxonomic richness of lakes for different periods of the Holocene, we highlight that our nutrient index is based on bedrock weathering, and the potential release of phosphorus, potassium, and calcium, which acts as a surrogate for alkalinity. During the Early Holocene, it is likely that nutrient release started immediately after deglaciation when liquid water was abundant (*39*) and light-demanding and disturbance-tolerant pioneer species could have survived on the nutrient-poor microhabitats, and thus showed weak overall association with the nutrient index. With continued warmer and possibly wetter conditions, leaching and nutrient release would have increased, thereby promoting richness in the Middle and Late Holocene. Relevant here is the fine spatial scale of the calcareous (alkaline) and siliceous (acidic) bedrock in northern Fennoscandia, with small and often linear, outcrops of metamorphic carbonate. This contrasts with the large calcareous limestone massifs found in younger geologies such as the European Alps, which have been shown to have effects on diversity over both short and long timescales (*11*). Given that there is also a positive association between nutrient index and total richness (representing a subset of the regional species pool), it is reasonable to consider nutrient index as an important driver for species pool development and hence regional richness (*40*). Indeed, it is the floristic variation between sites (beta-diversity) that accounts for the large difference between the local and regional species pools even today (*12*, *41*).

### A steep Early Holocene increase in plant richness

The highest rate of increase in richness and in local and regional species pools is observed in the Early Holocene 11.7-8.3 ka. Due to their significant correlation, we cannot distinguish the effect of dispersal lags from temperature, and both factors likely contributed to the observed increase in diversity. Climate was also the driver for deglaciation, which increased the area available for colonisation. Three of our records span a longer time period than examined here (Langfjordvannet: 16.7 ka, Nordvivatnet: 12.7 ka, Sandfjorddalen: 12.5 ka; Fig. S1F,H,I), and they, as well as macrofossils (14.5 ka, *42*) and pollen records (13.9 ka, *43*) from other coastal sites in northern Fennoscandia, show that an Arctic pioneer vegetation established towards the end of the Younger Dryas period (12.9-11.7 ka) and into the Early Holocene. Thus, a species pool already existed at least along the coast at the start of our study period, which is not the case for some of the inland sites (Gauptjern, Horntjernet, Kuutsjärvi) that were deglaciated after the onset of the Holocene. Nevertheless, all sites exhibit a strong increase in richness independent of location relative to deglaciation.

Especially during the rapid warming at 11.7-10 ka, we find a high increase in richness. Additional factors other than climate and availability of land may have influenced richness in this period. For example, biotic factors such as low levels of competition may have facilitated establishment (*44*), and abiotic factors, particularly paraglacial processes, may have produced disturbance at the local scale (*45*). On the other hand, dispersal lags may have limited richness and species pools, as for example the 400-year time lag between climate and arrival of birch woodland which was estimated based on plant macrofossils recovered from sediment cores (*46*). Nevertheless, the overall rapid increase in diversity in an early phase of colonization is also recorded in pollen studies (*19*), and expected given that they cover the development from pioneer to established vegetation communities.

Our richness patterns show a continued strong increase after ~11 ka, when the major expansion of birch forest took place, and after ~10 ka when pine expanded into the region (*21*). Thus, in contrast to the decrease in richness due to forest expansion observed in pollen studies (*13*, *19*), we found a general increase in richness through time. This may be because *sed*aDNA analyses are less sensitive to swamping by trees than pollen analyses and therefore better reflect habitat complexity (*25*–*27*).

### Middle Holocene richness continues to increase

The moderate increase in local and regional species pools during the Middle Holocene (8.3-4.2 ka) was marginally related to climate. The NGRIP record shows a temperature peak (end of the Holocene Thermal Maximum) followed by slight cooling during this period. Richness levelled off in only two lakes (Nordvivatnet and Sandfjorddalen) and one lake (Langfjordvannet) showed a hump in richness, which we assume is due to local factors. For Gauptjern, palynological richness fluctuates around eight taxa for this period (*47*), whereas our *sed*aDNA data show a clear increase. Pollen studies from northern Fennoscandia have shown contrasting patterns through this period, including stable levels of richness along a spruce-pine-birch tundra transect (*21*) and an overall increase in richness (*20*). The closest sites previously studied for *sed*aDNA show stable richness at Varanger in Finnmark (*35*), increasing richness in Svalbard (*36*), and fluctuating high richness in the Polar Urals (*34*). Seen from a regional perspective, our richness curves are similar to those found in the temperate zone of Europe, where a Middle Holocene richness increase is inferred to be due to human impact, but differ from those of the boreal zone (*19*), probably due to lower influence of Holocene tree expansion in the *sed*aDNA data. Thus, in contrast to many pollen studies, our *sed*aDNA data show an increase in richness and species pool for the Middle Holocene. As the climate was relatively stable during this period, we speculate that the increase in richness may have been mostly driven by dispersal lags, resulting in delayed establishment of some taxa in the region, and/or early habitat diversification allowing for a greater variety of niches, including the development of heathland, meadows, and mires (*14*).

### Late Holocene richness nears a plateau

The regional species pool clearly levelled off during the past few millennia suggesting that a near saturation point was reached. The slight cooling and well-known instability in this period (*48*) negatively impacted richness, potentially as a consequence of the cooling-induced withdrawal of the forest in the region (*14*, *21*). Palynological richness in northern Fennoscandia also increases slightly (*47*) or is variable (*21*) during this period. Richness also increases at sites in the boreal and temperate regions of Europe, mainly due to human land use impacts (*19*, *22*). The reason for levelling off at the regional scale in northern Fennoscandia is likely due to the near-saturation of the regional species pool and the overall low impact of human land use within the catchments.

In contrast to the regional scale, our data suggest that the local species pools and richness are not yet saturated. This is in contrast to what has been observed in studies of modern vegetation, where there appears to be no effect of time since glaciation for local (plot level) richness, whereas a legacy of the ice age is inferred for richness at the pan-Arctic (floristic region) scale (*16*). This apparent contradiction may be the result of scale and environmental spatial variation. Our catchments are larger than the plots studied by Stewart et al. (*16*), and therefore allow for co-existence of different vegetation types. Soils develop slowly on hard felsic and mafic rocks and have low buffering capacity resulting in nutrient loss and the partial development of oligotrophic vegetation types such as acid heaths and ombrotrophic mires. These have their own floras and some species are restricted to these environments. Indeed, mires and heath vegetation expanded in the region during the Late Holocene (*14*, *21*). Depending upon the local bedrock, a given area may thus gradually come to include additional vegetation types, allowing more hardy species to grow and total richness to increase while retaining the more demanding species in more favourable areas of the catchment. In addition, infilling of the lake creates wetland zones that also may include terrestrial taxa. Thus, a continued increase in richness and local species pools may be due to habitat diversification.

### Conclusions

We are aware that our study has several limitations. First, one should be aware that *sed*aDNA analyses based on p6-loop metabarcoding has taxonomic biases, as some species-rich families such as Salicaceae, Poaceae and Cyperaceae are poorly resolved due to haplotype sharing (*30*, *49*). We have used a local reference database to maximise taxonomic resolution and note that this is also an issue for traditional palynological proxies. Second, we acknowledge that the initial steep increase for our species pool estimates at the start of the Holocene constitute a sampling probability artifact, as plant taxa are known from the region in the late glacial (*42*, *43*). However, we note that a consistent steep increase continues throughout the Early Holocene interval, which we do not consider to be a sampling artifact. Third, we assume that the NGRIP record is representative of the climate of northern Fennoscandia. This record is in accordance with reconstructions of local climate in northernmost Fennoscandia based on macrofossils and pollen, although local variation does exist especially due to the proximity of the Norwegian Coastal Current, which is an extension of the Atlantic Gulf Stream (*43*). Lastly, we have not considered human impact as a potential driver of richness or species pools, but emphasise that this is considered to have been low in northern Fennoscandia as compared to other regions in Europe (*14*).

We present terrestrial plant richness and species pools inferred from 10 Holocene *sed*aDNA records covering environmental gradients in northern Fennoscandia. Our development of new quality control metrics to standardize data, together with improved taxonomic precision and known source areas (hydrological catchments of the lakes) allows for meaningful estimates of taxonomic richness, its spatial variation, and temporal trends, which show a unique increasing pattern of terrestrial plant richness over the Holocene. Our data reveal a steep increase in diversity in the Early Holocene related to the concurrent increase in temperature at that time and abundant vacant niches. However, the richness and the local and regional species pools continued to increase through the Middle and Late Holocene, although at a slower rate, suggesting that dispersal lags and habitat diversification had a major impact on diversity in these latter periods. In addition, we found that local nutrient levels, calculated based on bedrock type, had a strong impact on the overall levels of richness. Individual differences were observed among our sites, but our novel combined and standardized *sed*aDNA analyses of 10 sites provides a superior representation of the overall regional patterns in plant taxonomic richness over the Holocene as compared to traditional approaches. We suggest that plant diversity will continue to, and perhaps markedly, increase in northern Fennoscandia as a consequence of the northward movement of warm adapted species due to ongoing climate warming (sensu *1*). Expanding human impacts, including niche construction and introductions, within the region would further increase diversity. However, increases in diversity may be tempered by dispersal lags and the time taken for habitats to diversify as is suggested by our Holocene data set.

Integration of long-term paleo- and contemporary ecological data will be key to understanding, predicting, and managing the consequences of ongoing climate warming in northern ecosystems. Our study showcases how regional scale *sed*aDNA data can provide refined paleoecological insights into richness and species pools, as compared to traditional proxies, which will increase the precision of integrative ecological models.

## Materials and Methods

### Study area, site selection, and properties

The study area covers northernmost Fennoscandia above the Arctic Circle (at 67.75-70.43 N and 19.62-30.02 E) with nine lakes in Norway and one in Finland (Fig. 1, Data set S1). Site climatic and environmental properties are given below, whereas geology and vegetation descriptions are in the Supplementary Materials, SI1 and SI2.

#### Site selection

We selected 10 sites with the aim of minimising variation in environmental and potentially taphonomic variables (such as lake size and altitude), whilst covering the variation of biogeographical variables and particularly intra-regional climate. All sites are therefore small to medium sized lakes, with small catchments, no or minimal input streams, and a location above the maximum marine limit. We aimed to cover the major vegetation types in the region including the northernmost spruce and pine forest, the more widespread birch forest as well as subarctic grassland below the present-day treeline (Data set S1).

#### Climate

The Norwegian sites are located in the three most northerly Norwegian temperature regions (TR4-TR6) and precipitation regions (RR11-RR13), as defined by Hansen-Bauer and Nordli (*50*). Statistical analysis has established that these are the most homogenous sub-regions related to common drivers of regional forcing, particularly sea-level pressure over the North Atlantic and North Atlantic Oscillation (NAO) (*51*). The Finnish site (Kuutsjärvi) is close to the easternmost part of Norway, with a similar continental climate.

#### Environment

The site environmental data was taken from published sources: site altitude, lake area, catchment area and climatic variables was generated from the NAVINA-NVE online tools (https://www.nve.no/karttjenester/) and catchment geology from the Geological Survey of Norway (NGU) online mapping system (https://www.ngu.no/en). Given that the role of parent material is particularly important for plant growth in both sub-arctic and Alpine environments (*52*) it was deemed potentially valuable to classify the sites by the geologically derived nutrient resource environment (see Equation 2). In most cases values from the bedrocks of the sites are not available so typical values measured from a suite of rocks from the Caledonian-nappe metamorphic rocks from north Troms was used (*12*) the values of which can be found in Data set S1. However, in the case of Nordvivatnet and Sandfjorddalen values could be taken from studies of the alteration of Neoproterozoic tillites of the Varanger region (*53*). The major methodological uncertainties with such an index is the role of both the moraine cover and the development of a soil organic matter store. The moraine cover is typically pre-weathered to some extent and probably reduces the effect of the local bedrock. This approach is appropriate for the geologically mediated plant nutrients and nutrient storage and cycling governed by the development of a soil organic matter (SOM) store.

### Fieldwork and lake sediment coring

We collected cores from seven of the ten lakes from northern Fennoscandia between February and April 2017 (Fig. 1; Data set S1), whereas cores from three lakes were obtained from previous studies (see below). For each of six of the lakes, we retrieved sediment in a single 10-cm diameter polyvinyl chloride (PVC) pipe, using a modified Nesje piston-corer (*54*) from the deepest and generally central part of the lake. At Lake Gauptjern, two Nesje cores were retrieved (EG03, EG13). We monitored for potential cross contamination using a DNA tracer mixed into Vaseline, which we applied to the piston immediately prior to coring. This tracer consisted of DNA extracted from a non-native plant, the Christmas cactus (*Schlumbergera truncata*) (see Supplementary Materials, SI3). After coring, we cut the pipe into 1.0-1.2 m long sections in the field and immediately sealed the ends to minimize contamination from modern environmental DNA (eDNA). For six of the lakes, we collected up to 1-m long cores using a 4-cm diameter rod-operated Multisampler (Eijkelkamp 12.42; Giesbeek, The Netherlands), which usually captured the water-sediment interface (Data set S2). At Sandfjorddalen, two sequential Multisampler cores were taken from the same hole, resulting in a 149 cm core. All core sections were kept cold during transport to and storage in a dedicated cold room (4 °C). During storage, water from the majority of the Multisampler cores leaked out, which resulted in shrinkage of these cores.

### Core sampling

The Nesje cores were split longitudinally, with one half used for destructive analyses (loss-on-ignition, LOI; *sed*aDNA) and the other for high-resolution imagery. Sampling for destructive analyses was conducted in the dedicated ancient DNA laboratory facility at The Arctic University Museum of Norway (TMU), using sterile implements. We sampled all cores starting at the base and fully cleaned the laboratory between sampling cores from different lakes. At each selected 1-cm layer, we first removed and discarded the top ~2 mm of exposed sediment surface. We then removed a further ~3 mm (~5-10 g) of the underlying sediment surface, which we retained for LOI analysis. Finally, we then sampled ~10-20 g of the remaining underlying sediment for *sed*aDNA analysis, taking care not to sample sediment immediately adjacent to the PVC pipe. To detect contamination from exogenous sources of environmental DNA, we took sampling negative controls that consisted of 100 μL of exposed molecular biology grade water.

For the Multisampler cores with remaining water and highly liquid surface sediment, we first siphoned off and discarded the water. We then siphoned and retained the liquid surface sediment at 1 cm intervals. For the remaining sediment, and for Multisampler cores that had shrunk due to water loss, we extruded the cores at 1 cm intervals, starting at the base of the core. At each interval, the outer ~5 mm of sediment was removed and retained for LOI analysis. The remaining ~3 cm diameter sediment was retained for *sed*aDNA analysis. Sampling conditions and controls were as described above.

We sampled sediment from three cores that had been previously collected and were available to us. These consisted of records from Langfjordvannet (*31*), Jøkelvatnet (*55*), and Kuutsjärvi (*56*). For the Langfjordvannet and Jøkelvatnet cores, sampling was conducted within the clean labs of GeoMicrobiology at the Department of Earth Science, University of Bergen, Norway following the approach mentioned above. Sampling of the Kuutsjärvi core was conducted in the Physical Geography department at Stockholm University, Sweden using sterilized tools and with all surfaces cleaned with bleach. We took negative sampling controls at both institutions as described above.

### Core photography and loss-on-ignition analyses

The intact core halves were photographed at high resolution using a Jai L-107CC 3 CCD RGB Line Scan Camera mounted to an Avaatech XRF core scanner at the Department of Geosciences, The Arctic University of Norway in Tromsø. We calculated mass LOI, by first drying samples 105 °C overnight and then igniting the sample at 550 °C for 2 hours. We report the percentage of dry mass lost after ignition (*57*). LOI data and high resolution core scanning imagery are presented in Data set S3 and Fig. S1.

### Composite core construction and age-depth modelling

We opportunistically collected macrofossils for radiocarbon (^14^C) dating during sampling for LOI and *sed*aDNA, where possible. If additional macrofossils were required, we sieved sediment to concentrate macroscopic plant remains suitable for dating. Two sieves of 500 and 250 μm were stacked while sieving to catch plant macrofossil remains. Ultrapure water from the Milli-Q system was used for sieving and cleaning, and samples were kept cool in Eppendorf tubes with water before shipping for dating. We photographed and identified all macrofossils prior to their destruction during radiocarbon dating. Samples were radiocarbon dated using Accelerator Mass Spectrometry (AMS) at the Poznań Radiocarbon Laboratory of the Adam Mickiewicz University, Poland (*58*) (Data set S3).

For multiple-core records from the same site, we aligned cores based on combinations of LOI values, visible stratigraphy, and/or radiocarbon dates to create composite core records (Fig. S1; Data sets S2, S3). For Gauptjern, we used the LOI profile and radiocarbon dates produced by Jensen & Vorren (*47*) to guide composite age-depth model construction (Fig. S1B; Data set S3). All reported depths are based on the composite cores and begin at the water-sediment interface, which was determined either by its successful capture, field notes, or previously published information (Data set S2). We note that the composite depths reported here differ for two of three previously published records (*31*, *47*, *55*) (Data set S3). For Langfjordvannet, we increased depths by 26 cm to account for the amount of sediment reported missing from the top (*31*), whereas for Gauptjern, we removed 340 cm (water depth) and adjusted remaining depths for differing deposition rates among all cores, due to an uneven bedrock surface (*47*).

We constructed Bayesian age-depth models using *bacon* v.2.3.4 (*59*) in R v3.4.4 (*60*), using the IntCal13 calibration curve (*61*). Each basal modeled age was ≤2 cm below the basal radiocarbon date, with the exception of Langfjordvannet where the basal radiocarbon date falls within a slump that extends to the base of the core. We were unable to confidently model the basal 31 cm of Jøkelvatnet and 22 cm of Kuutsjärvi, as extrapolated ages were highly influenced by accumulation rate priors. We fixed the top of each record to zero, based on the composite cores beginning at the water-sediment interface. The default section thickness of 5 cm was used for all age-depth models, with the exception of Sandfjorddalen, which shrank from the 149 cm collected to 92 cm analyzed. We therefore selected a 2 cm thickness for the Sandfjorddalen age-depth model. We excluded two dates (Poz-108675, Poz-108983) from the Sierravannet age-depth model that occurred in a putative flood layer (Fig. S1J).

### Sedimentary ancient DNA data generation

We performed all pre-PCR steps at the dedicated ancient DNA facilities at TMU, which are in a separate building to post-PCR facilities. We homogenized Holocene-aged DNA samples by holding samples on a pulse vortexer for ~1 min. We extracted DNA from 0.25-0.35 g of sediment (Data set S4) using a modified form of the Qiagen DNeasy PowerSoil PowerLyzer (Qiagen Norge, Oslo, Norway) protocol, following the Zimmermann *et al*. (*62*) protocol, as modified by Alsos *et al*. (*63*). We included one negative extraction control, consisting of no input, for every 10 sediment extractions. We also extracted DNA from 10 samples using the protocol 15 of Heintzman *et al*. (in prep) and six samples using the carbonate protocol of Capo *et al*. (submitted).

We amplified DNA and control extracts using ‘gh’ primers (*64*) that target the vascular plant trnL p6-loop locus of the chloroplast genome (Data set S5). The gh primers were unique dual-tagged with an 8 or 9 base pair tag, modified from Taberlet et al. (*65*). We used differing tag lengths to ensure that nucleotide complexity was maintained during amplicon sequencing runs. A total of eight PCR replicates were amplified per extract. We included negative PCR controls, consisting of water as input, to monitor for contamination during the PCR. We additionally included negative and positive PCR controls in the post-PCR lab, the latter of which consisted of one of six synthetic sequences (available at https://github.com/pheintzman/metabarcoding) (see also Supplementary Materials, SI4). These post-PCR lab controls were added to wells without disturbing other sealed sample and control wells and were used to monitor PCR reaction success. However, they are not comparable to other negative controls and samples, due to exposure to the post-PCR lab atmosphere, and so they were excluded from further analysis. We checked for successful amplification using gel electrophoresis (2% agarose).

We pooled up to 384 PCR products (the maximum number of available tags) and then cleaned the resulting pool following Clarke *et al*. (*35*). Each amplicon pool was then converted into a DNA library at either Tromsø or FASTERIS, SA (Switzerland). The Tromsø protocol used the Illumina TruSeq DNA PCR-free protocol (Illumina, Inc, CA, USA) with unique dual-indexes, except that the magnetic bead cleanup steps were modified to retain short amplicons, whereas FASTERIS used the PCR-free MetaFast protocol to produce single-indexed libraries following Clarke *et al*. (*35*). Each library was sequenced on ~10% of 2x 150 cycle mid-output flow cells on the Illumina NextSeq platform at either FASTERIS or the Genomics Support Centre Tromsø (GSCT) at The Arctic University of Norway in Tromsø, or on 50% of a 2x 150 cycle flow cell on the Illumina MiSeq platform at FASTERIS. Full sample preparation metadata is provided in Data set S5.

### Bioinformatics

We followed a bioinformatics pipeline that uses a combination of the ObiTools software package (*66*) and custom R scripts (available at https://github.com/Y-Lammers/MergeAndFilter). Briefly, we merged and adapter-trimmed the paired-end reads with SeqPrep (https://github.com/jstjohn/SeqPrep/releases, v1.2). We then demultiplexed the merged data using an 8 bp tag-PCR replicate lookup identifier (provided in Data set S5), which ignored the non-informative terminal base for 9 bp tags, and collapsed identical sequences. We removed putative artifactual sequences from our data, which may have derived from Illumina library index-swaps or PCR/sequencing errors. For each PCR replicate, we removed sequences represented by ≤2 reads. We next identified barcode sequences that had 100% identity agreement with a local taxonomic reference database (*ArctBorBryo*) containing 2445 sequences of 815 arctic and 835 boreal vascular plants, as well as 455 bryophytes (*30*, *49*, *67*). In addition, we matched our data set to the *EMBL* (rl133) nucleotide reference database. We separately compared our barcode data set against the barcode sequence of the DNA tracer, with the closest match consisting of 85% identity (see Supplementary Materials, SI3). We therefore consider the DNA tracer not to be present in our data set. We further removed identified sequences that 100% matched against two *blacklists* (https://github.com/Y-Lammers/Metabarcoding_Blacklists) consisting of either synthetic sequences (n=6), sequences that represented homopolymer variants of a more read-dominant sequence, a potential random match, or food contaminants (n=111) (Data set S6). We further removed any sequences represented by fewer than 10 reads and/or three PCR replicates within the entire data set, as well as 61 low-frequency sequences that were only retained by analysis of the entire data set but removed if analyses were conducted on a per-lake basis (Data set S6). If multiple sequences were assigned to the same taxon, then the data were merged using the sum of all assigned reads and the maximum number of PCR replicates (Data set S7). The final taxonomic assignment of the retained sequences was determined using regional botanical taxonomic expertise by Alsos and following the taxonomy of the Panarctic Flora (*68*) and Lid’s Norsk Flora (*69*). We identified two species of *Vaccinium* based on a poly-A region at the 3’ end of the p6-loop locus. If there were ≤5 or >8 As, then barcodes were respectively assigned to *V. myrtillus* or *V. vitis-idaea*. We further excluded barcodes that were identified above the family level based on alignment to the local reference database, *ArctBorBryo*. Among the identified plant taxa, only terrestrial vascular plants and bryophytes were retained for all downstream analyses (Data set S6). We only included Holocene-aged (11,700 ka to present) samples for downstream analysis. We note that we use the term taxonomic richness to include taxa identified to various ranks from the species to family levels.

### Assessment of sedimentary ancient DNA data quality

Our *MTQ* and *MAQ* score thresholds excluded all negative controls, which had an *MTQ* of <0.75 and *MAQ* of <0.1. Across our entire data set, 16 samples were extracted more than once. We included data from the DNA extract that yielded the greatest *MAQ* score. In three cases with equal *MAQ* scores, we selected replicate one for inclusion (Data set S4).

After data filtering, we found that there was often large variation in the counts of retained reads between PCR replicates within a sample (from hundreds to tens of thousands). Although read-dominant barcodes are likely to be detected in all PCR replicates, there is likely to be dropout of other barcodes in replicates with lower counts of retained reads. In contrast, rare barcodes are more likely to be detected in replicates with high retained read counts. We therefore developed a barcode detectability measure - *wtRep* - to account for differences in relative counts of retained reads, by weighting PCR replicates based on retained read count relative to the total retained read count across all PCR replicates, on a per sample basis (see Equation 3). For example, if a barcode were detected in replicates one and three, but undetected in the remaining six replicates, the *wtRep* would as shown in Equation 1.

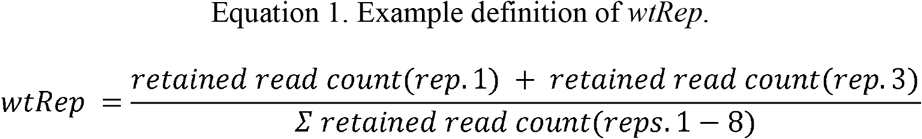

If a PCR replicate were not represented in the retained read data, then it would not contribute to the *wtRep* score. A limitation of the *wtRep* score is that it will overrepresent detections in samples or negative controls with few barcodes and/or detections. For this reason, we only applied *wtRep* for a sample if the average proportion of replicates across the sample was ≥0.33 and there were ≥10 barcodes present after filtering. For samples that failed this threshold, we used a standard proportion of PCR replicates as a measure of detectability (e.g. 0.25 for the above example).

We further explored the quality of our *sed*aDNA data by examining four measures. For each sample, we calculated the (1) total count of raw reads (summed across PCR replicates), (2) mean barcode length (in base pairs, bp) across all retained barcodes, (3) mean proportion of weighted PCR replicates (*wtRep*; see above) across all final barcodes, and (4) proportion of raw reads assigned to terrestrial plant taxa. We compared each of these four measures to both observed taxonomic richness (Hill-N0) and/or time.

### Numerical and statistical analyses

Using the proportion of weighted PCR replicates (*wtRep*), we measured taxonomic richness (diversity) based on Hill numbers (N0 and N1) (*70*), as they are easily interpretable and provide information on both the rare and common taxa within a community (*13*). Hill numbers have been widely used as common metrics to link different ecological attributes (*71*) as well as increasingly used in DNA-based diversity analyses (*72*). For each sample, we calculated taxonomic richness as Hill N0 (total number of observed taxa), and number of abundant taxa as Hill N1 (see Equation 2), which is the exponent of the Shannon index (*13*). To evaluate if the disproportionate sequencing depth has affected the taxonomic richness estimates, we also calculated rarefied taxonomic richness based on the lowest number of reads assigned to a sample within a lake. We calculated Pearson’s product-moment correlation between rearefied and observed (Hill N0) taxonomic richness to evaluate the correspondence between approaches (Table S2).

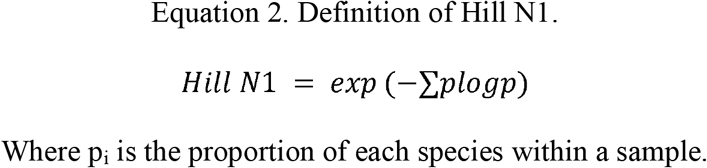

We used generalized additive models (GAMs) (*73*) to evaluate temporal biodiversity changes during the Holocene. GAMs are very efficient at uncovering nonlinear covariate effects (*74*) and handling non-normal data that are typical in palaeoecology (*75*). We treated Hill N0 and N1 as the response, and median calibrated age of the samples as predictor variables, and used the “poisson” family with log link. The Hill N1 was rounded to the nearest whole number prior to GAM analysis. To account for residual temporal autocorrelation between samples, we also included a continuous time first-order autoregressive process (CAR(1)) in generalized additive mixed models (GAMM; (*75*)). For both GAM and GAMM models, the fitted lines are based on the predicted values for 300 points covering the entire range of sample age for each lake. We used a critical value from the t distribution to generate a pointwise 95% confidence interval (*75*). We found near identical results for taxonomic richness between GAM and GAMM models (Fig. S7; Table S3). In the cases of two shorter cores from Nesservatnet and Sierrvatnet, the GAMM provided a reasonable fit to the data, and hence was included in the main results.

We also evaluated how the local and regional species pools affected richness estimates at respective scales. We define the local and regional species pool as the cumulative number of taxa recorded within a lake or the total region, respectively. First, we calculated the cumulative number of detected taxa from the oldest to the youngest samples across all lakes as an indicator of the development of the regional species pool, and compared that qualitatively to the taxonomic richness of all samples through time. In addition, we generated a regional species pool of total species recorded within each 500-year time bin. We extracted major trends of richness based on 500-year bins using GAM for the Early-, Middle-, and Late Holocene. We also used GAM to highlight the regional trend in taxonomic richness through time. Then, we performed linear regression by considering the mean number of taxa within a lake, and within a 500-year time bin as the response variables, and the respective species pools as the predictor variables to test whether observed richness is correlated to the species pools of respective scales.

To examine the relationship of climate and diversity estimates, we used oxygen isotope (δ^18^O) values from the North Greenland Ice Core Project (NGRIP) (Andersen *et al.*, 2004) as a proxy for temperature. This has limitations as a regional record for northern Fennoscandia because of geographic distance, but its advantage is that it is independent of vegetation-based reconstructions and it covers the whole period of interest. It shows similar trends in annual temperature as both proxy-based and simulated reconstructions for the northern North Atlantic region (Marsicek *et al.*, 2018). Because summer temperature and growing degree days (GDD) are strong drivers of vegetation response to climate in the north, we note that Holocene GDD sums probably remained higher than present through much of the Holocene, as patterns of both seasonality and season length changed (Marsicek *et al.*, 2018). The Early Holocene temperature change was steeper than the Middle- and Late Holocene, and we expect the richness pattern to differ among those periods. Thus, we assigned δ^18^O data of 50 years resolution to the nearest age estimate of samples, and split diversity estimates and δ^18^O values into three periods (Early:11.7-8.3 ka, Middle: 8.3-4.25 ka, and Late Holocene: 4.25-0.0 ka) following Walker *et al*. (*76*). To evaluate how changes in temperature affected the richness pattern of the different Holocene periods, we compared regression slopes of the Middle and the Late Holocene to the Early Holocene using a linear mixed model with taxonomic richness as the response and an interaction between δ^18^O and the Holocene period as predictor along with lakes as the random variable.

We used a new semi-quantitative nutrient index derived from the sum of the phosphorus (P), potassium (K), and calcium (Ca) content of the rocks modified by a measure of weatherability, in this case the extended Moh’s hardness of the least resistant major mineral in the rock type (H_min_) (see Equation 3). The natural logarithm of Ca content was used as this has been shown to have a strong relationship with pH, which is critical to the availability of nutrients especially P (*12*). We performed linear regression treating mean taxonomic richness of different periods of the Holocene as the response and nutrient index as the predictor. The richness data were log transformed while evaluating the impact of temperature and nutrients.

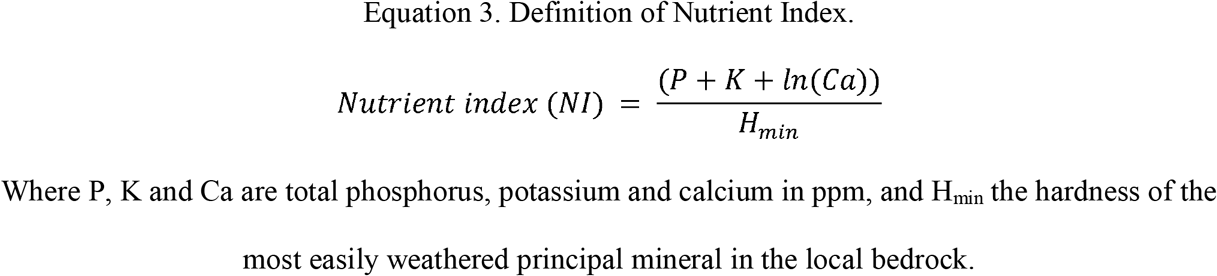

Where P, K and Ca are total phosphorus, potassium and calcium in ppm, and H_min_ the hardness of the most easily weathered principal mineral in the local bedrock.

Unless otherwise stated, all the analyses were performed using the *vegan* package (*77*) in R and base *R (60)*. The library *mgcv* (*73*) was used for GAM model building. All plots were created using *ggplot2* (*78*).

## Supplementary Materials

Supplementary Information 1. Site geology.

Supplementary Information 2. Site vegetation.

Supplementary Information 3. DNA tracer.

Supplementary Information 4. Positive control synthetic sequences.

Fig. S1. Alignments of core LOI, high-res. imagery, and Bayesian age-depth models.

Fig. S2. A potential flood event does not impact the Sierravannet diversity trend.

Fig. S3. Observed taxonomic richness in each sample by lake and time including samples not passing quality controls.

Fig. S4. Six measures of *sed*aDNA data quality by lake and time.

Fig. S5. The assignments of reads processed by the bioinformatic pipeline.

Fig. S6. The accumulated regional species pool and taxonomic richness of each sample across the Holocene of northern Fennoscandia excluding two temporally-short records from Nesservatnet (EG02) and Sierravannet (EG07).

Fig. S7. Comparison between GAM and GAMM(CAR(1)) models of taxonomic richness through time.

Table S1. Summary of all data used or generated in this study.

Table S2. Correlations between observed and rarefied taxonomic richness for each lake.

Table S3. Summary of generalized additive models (GAMs), and generalized additive mixed models (GAMMs) with a continuous time first-order autoregressive (CAR(1)) process.

Data set S1. Geographic and site metadata for the ten lakes.

Data set S2. Composite core construction and Bayesian age-depth modelling.

Data set S3. Sample metadata, including depths, LOI values, dates, and modelled ages.

Data set S4. Full sample metadata including QC and bioinformatic sequence processing.

Data set S5. Primer tag to sample lookup, library preparation, and sequence accession data.

Data set S6. List of all identified barcodes, including those blacklisted, and their taxonomic assignments and functional groups.

Data set S7. Read counts and PCR replicate detections for all retained taxa across all samples.

## General

We thank Roseanna Mayfield and Kristian Sirkka for assistance with fieldwork; Marie Føreid Merkel, Sandra Garces Pastor, and Yann Belov for technical assistance in the lab; Karina Monsen for assistance with high-resolution photography; Eivind Støren, Department of Earth Science, University of Bergen for providing access to cold rooms and helping to locate and transport cores; Ruth Paulssen and the Genomics Support Centre Tromsø (GSCT) at UiT-The Arctic University of Norway for amplicon sequencing; Dorothee Ehrich for fruitful discussions and coring at Sandfjorddalen; Matthias Forwick for help organizing scanning of cores; and Lasse Topstad and Leif Einar Støvern for providing photographs. DPR is thankful to Keshav Prasad Paudel for his help in map creation. Bioinformatic analyses were performed on resources provided by UNINETT Sigma2 - the National Infrastructure for High Performance Computing and Data Storage in Norway.

## Funding

The study is part of the project “ECOGEN - Ecosystem change and species persistence over time: a genome-based approach”, which is financed by Research Council of Norway grant number 250963/F20.

## Author contributions

IGA, KAB, NGY, TA, and MEE designed the research and raised the funding; IGA, AGB, DPR, PDH, FJAM, YL, KAB and KEL did the fieldwork; JB, KFH, and JSS provided resources; DPR, PDH, KEL and IP did the laboratory work with input from IGA and AGB; TG performed radiocarbon dating; PDH built composite cores and performed age-depth modelling with input from AGB, TG and KFH; YL and PDH designed the bioinformatic pipeline with input from DPR and IGA; IGA and DPR verified and curated barcode taxonomic assignments; PDH and YL designed and performed the quality control checks with input from DPR and IGA; DPR did the statistical analyses with input from NGY and KAB; YL, DPR and PDH curated the data; DPR, PDH, AGB and IGA drafted the manuscript; and all authors have reviewed and approved the final manuscript.

## Competing interests

The authors declare no competing interests.

## Data and materials availability

Raw sequence data have been deposited in the European Nucleotide Archive (ENA) at project accession PRJEB39329, with sample accessions ERS4812035-ERS4812048. All radiocarbon, loss-on-ignition, and processed sedaDNA data are available in the Supplementary Materials. Pre-filtered ObiTools tsv output files have been uploaded to figshare (DOI: [available upon acceptance]). Scripts are on Github with URLs cited in the Materials and Methods section.

## Notes

### Competing Interest Statement

The authors have declared no competing interest.

### Summary of Updates

Introduction, results, and discussion shortened and sharpened. Figure 2 is new, Figure 3-7 revised. Conclusion section extended and improved. Supplementary files updated.

