## Supplementary Materials for "Sedimentary ancient DNA shows terrestrial plant richness continuously increased over the Holocene in northern Fennoscandia"

**SI.1. Site geology**.

The sites are split between the Precambrian (Neoproterozoic) province of Varanger (Nordvivatnet, Sandfjorddalen), the Caledonian metamorphic nappes of the western mainland and islands (Nesservatnet, Gauptjern, Sierravannet, Eaštorjávri South, Landfjordvannet, Jøkelvatnet) and the Proterozoic basement of southern Finnmark and northern Finland (Horntjernet, Kuutsjärvi). Lithologically they are divided between felsic rocks (gneiss, granites, and quartzites) and mafic rocks (amphibolites, hornblende, gabbro) and one lake on carbonate rocks (marble). However, locally there are variations which ameliorate the bedrock chemistry such as the presence of dolomitic clasts in Neoproterozoic conglomerates (Nordvivatnet), variable mica contents and Quaternary sediments including beach barrier deposits (Nordvivatnet), fluvioglacial gravels (Sierravannet) and moraines of variable thickness (see Data set S1). The soils of the catchment are dominated by relatively thin soils, with lithomorphic soils (leptosols FAO, 1998) on bedrock and moraines becoming gleyic, humic and podzolic locally. Only one catchment has any glacial ice in the catchment (Jøkelvatnet) and none are on permafrost, although Kuutsjärvi is marginal to the zone of discontinuous permafrost (*81*). Most of the lakes are glacial in origin although two are kettle hole lakes (Sierravannet, Kuutsjärvi) and they have variable dates of deglaciation with 5 from before GS-1 (11.8 ka) and 5 from after the Younger Dryas (c. 11.6-10.6 ka).

**SI.2. Site vegetation**.

The vegetation description in Data set S1 is based on unpublished field observations by Alsos, I.G., Bråthen, K.A., Rijal, D.P., and Lorberau, K. A more detailed description of the modern-day vegetation is presented below here:

Eaštorjávri South (EG11) make up a mostly treeless sub-arctic grassland with scattered individuals of mountain birch and *Salix* thickets close to the lake. Forbs present include *Alchemilla alpina, Angelica archangelica,* *Omalotheca norvegica, Omalotheca supina, Oxyria digyna, Pedicularis lapponicum, Rhodiola rosea, Solidago virgaurea, Taraxacum officinale, Trientalis europaea, Veronica alpina* and *Viola biflora.* There are some *Betula pubescens* trees and shrub *Salix* spp*.* but the woody species consist mainly of the dwarf shrubs *Betula nana, Empetrum nigrum, Salix herbacea, Vaccinium myrtillus, Vaccinium uliginosum* and *Vaccinium vitis-idea*. Graminoids include *Anthoxanthum odoratum, Avenella flexuosa, Calamagrostis sp, Deschampsia* spp*., Carex* spp*., Festuca ovina, Juncus trifida, Phleum alpinum* and *Trisetum spicatum*.

Gauptjern (EG03/13) is situated at the upper limit of pine in the region, and the vegetation is dominated by a rich birch forest with scattered pines. The field layer is dominated by dwarf shrubs *Empetrum nigrum, Vaccinium myrtillus, Chamaepericlymenum suecicum* and *Vaccinium vitis-idea* in addition to the tall forbs *Cicerbita alpina, Filipendula ulmaria*, *Trollius europaeus* and *Geranium sylvaticum* and the small forbs *Bartsia alpina*, *Geum rivale*, *Ranunculus* spp.*, Rumex acetosa* and *Solidago virgaurea*. Grasses include *Deschampsia cespitosa, Avenella flexuosa*, and *Anthoxanthum odoratum*. There are also the ferns *Dryopteris* spp. and *Athyrium* spp*.* See full species list for this site in Alsos *et al*. (*28*).

Horntjernet (EG05) lies within the northernmost continuous pine forest, and the surrounding vegetation is dense pine forest with some areas of mire. The forest is dominated with *Pinus sylvestris* trees and secondarily *Betula pubescens* with an understory of the dwarf shrubs *Betula nana, Empetrum nigrum, Calluna vulgaris, Vaccinium myrtillus, Vaccinium vitis-idaea, Rubus chamaemorus* and *Rhododendron tomentosum* along with the grass *Avenella flexuosa*. The mire areas contain the same dwarf shrub species with in addition the graminoids *Eriophorum vaginatum, Trichophorum cespitosum*, and *Carex* spp.

Jøkelvatnet (EG17) is in the alpine zone and no trees were recorded within the catchment. There are mire areas dominated by *Carex* spp*.* with *Eriophorum* spp. and *Betula nana,* and alpine heath dominated by dwarf shrubs *Empetrum nigrum, Vaccinium uliginosum, Vaccinium vitis-idea, Arctous alpinus*, and *Calluna vulgaris* along with the graminoids *Avenella flexuosa*, *Juncus trifidus* and *Festuca ovina*.

Kuutsjärvi (EG21) is in the northernmost spruce forest of the region, and the forest is dominated by *Picea abies,* with some *Pinus sylvestris* and *Betula pubescens.* The field layer is dominated by *Vaccinium myrtillus* and *Empetrum nigrum*. Heath areas also include *Arctostaphylos alpina* and *Antennaria dioica*.

Landfjordvannet (EG15) has vegetation around it today consisting of a rich and open mountain birch forest with an understory of the forbs *Cicerbita alpina, Filipendula ulmaria, Geranium sylvaticum* and *Aconitum lycoctonum*, the graminoids *Agrostis capillaris* and *Deschampsia cespitosa* as well as the fern *Athyrium filix-femina*. There are also areas of treeless heath dominated by the dwarf shrubs *Betula nana, Empetrum nigrum, Vaccinium myrtillus* and *Chamaepericlymenum suecicum* along with the grass *Avenella flexuosa*.

Nesservatnet (EG02) is at the northwestern limit of pine, and the vegetation is dominated by open mountain birch forest with some pine and heath and mire vegetation. The dominant species are *Betula pubescens* with *Betula nana, Empetrum nigrum, Calluna vulgaris, Vaccinium uliginosum* and *Vaccinium vitis-idaea*, and with *Rubus chamaemorus* and *Oxycoccus microcarpus* in the wetter areas. Also present are the graminoids *Avenella flexuosa, Festuca ovina, Carex* spp*.,* *Eriophorum vaginatum* and *Trisetum spicatum*.

Nordvivatnet (EG10) is surrounded by a mixture of mostly *Betula pubescens* forest with smaller wetland areas and species rich meadows. The field layer is dominated by dwarf shrub *Empetrum nigrum* as well as the grass *Avenella flexuosa*. Towards the west side there is treeless heath dominated by the dwarf shrub *Empetrum nigrum* with *Betula nana, Juniperus communis, Chamaepericlymenum suecicum, Vaccinium uliginosum,* and *Vaccinium vitis-idea,* in addition to the grasses *Avenella flexuosa* and *Anthoxanthum odoratum*, the forbs *Solidago virgaurea*, *Pedicularis lapponica*, and *Trientalis europaea*, and the fern *Gymnocarpium dryopteris*.

Sandfjorddalen lake (MS06) is today surrounded by treeless arctic-alpine heath and mire vegetation with some shrub thickets. The heath is dominated by the dwarf shrubs *Betula nana* and *Empetrum nigrum* with *Vaccinium myrtillus, Phyllodoce caerulea, Chamaepericlymenum suecicum* and *Salix herbacea*. The subdominant vegetation consists of the graminoids *Avenella flexuosa, Agrostis mertensii, Luzula multiflora* and *Anthoxanthum odoratum* as well as the forbs *Solidago virgaurea* and *Rumex acetosa*.

Sierravannet (EG07) also lies within the northernmost continuous pine forest. It is surrounded by a planted pine forest with some larch and a native birch forest on either side of a wet swamp area. Both the forest dominated by *Pinus sylvestris* and that with *Betula pubescens* have an understory of *Vaccinium myrtillus, Avenella flexuosa*, and *Empetrum nigrum*. There are also portions of the birch forest with the shrub *Juniperus communis* and with the ferns *Gymnocarpium dryopteris* and *Dryopteris expansa*. Subdominant species include the forbs *Solidago virgaurea, Potentilla norvegica, Trientalis europaea,* and *Melampyrum pratense*.

**SI.3. DNA tracer**.

We generated a trnL p6-loop barcode sequence from the same individual used to generate our DNA tracer, a Christmas cactus (*Schlumbergera truncata*) located at The Arctic University Museum of Norway. The barcode was generated via shotgun sequencing of the source plant, according to the methods described in Alsos *et al*. (*82*). We extracted the p6-loop sequence from an assembled chloroplast assembly with the ecoPCR package (v0.2: Ficetola *et al*. (*83*)). We compared all barcodes generated from our *sed*aDNA records against the tracer barcode sequence, using the ecoTag function from the ObiTools software package, but the tracer barcode was not present in our lake sediment data set. The *sed*aDNA-derived barcode that most closely resembled the tracer sequence was removed during data filtering and had a 100% identity match to the cactus *Selenicereus atropilosus*, and matched the tracer sequence with only 85% identity. The *S. atropilosus*-identified barcode was detected in three different samples; once in Horntjernet (one PCR replicate with four reads) and in two different samples from Langfjordvannet (one PCR replicate each with eight and seven reads, respectively). This detection of *S. atropilosus* is therefore unrelated to the DNA tracer, as the Langfjordvannet core was sampled in Bergen in the absence of tracer.

>DNA_tracer_p6-loop_barcode Schlumbergera truncata p6loop; taxid=3595;

CTCCTTTTTTTTTTTGAAAAAAAAAAGCAAAAAATAAGGGTTCAGAAAGCAAGAATAAAAAAAAAG

**SI.4. Positive control synthetic sequences**.

We generated six novel synthetic oligos, consisting of a randomized 30 or 60 bp sequence flanked by the *trn*L p6-loop gh primer binding sites, to act as positive PCR controls in our data set. We note that our first batch of tagged gh primers, which were used to generate the data in sequencing run JIE (Data sets S4, S5), were manufactured one month after, but in the same facility, as the synthetic oligos. We noticed that there was trace contamination of the synthetic sequences in these primer sets, and consequently synthetic sequences are present in samples and negative controls from run JIE. The pattern of which synthetic sequence(s) were associated with which tagged primer set, coupled with the fact that the synthetic oligos were never taken to the pre-PCR facility and that independently produced primers did not show this issue in comparative experiments using the same PCR master mix, preclude this being a contamination event in our lab. After ordering a new batch of tagged gh primers, generated in a separate facility, we did not observe this issue in any other subsequent runs (Data set S4). We highlight this experience as a warning to researchers considering the manufacture of positive control sequences, and to demonstrate that primers can be a source of PCR reagent contamination.

**A**

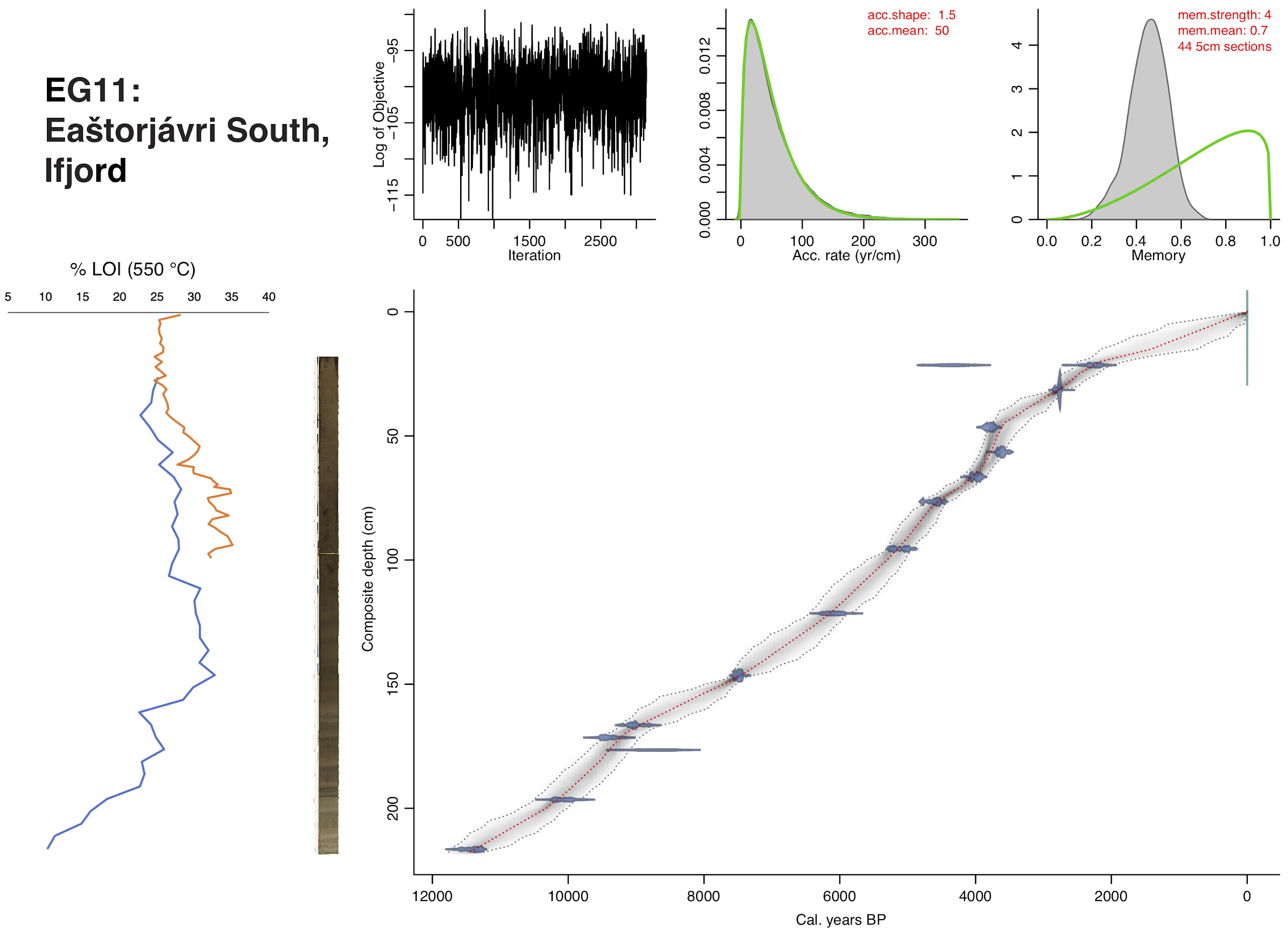

**B**

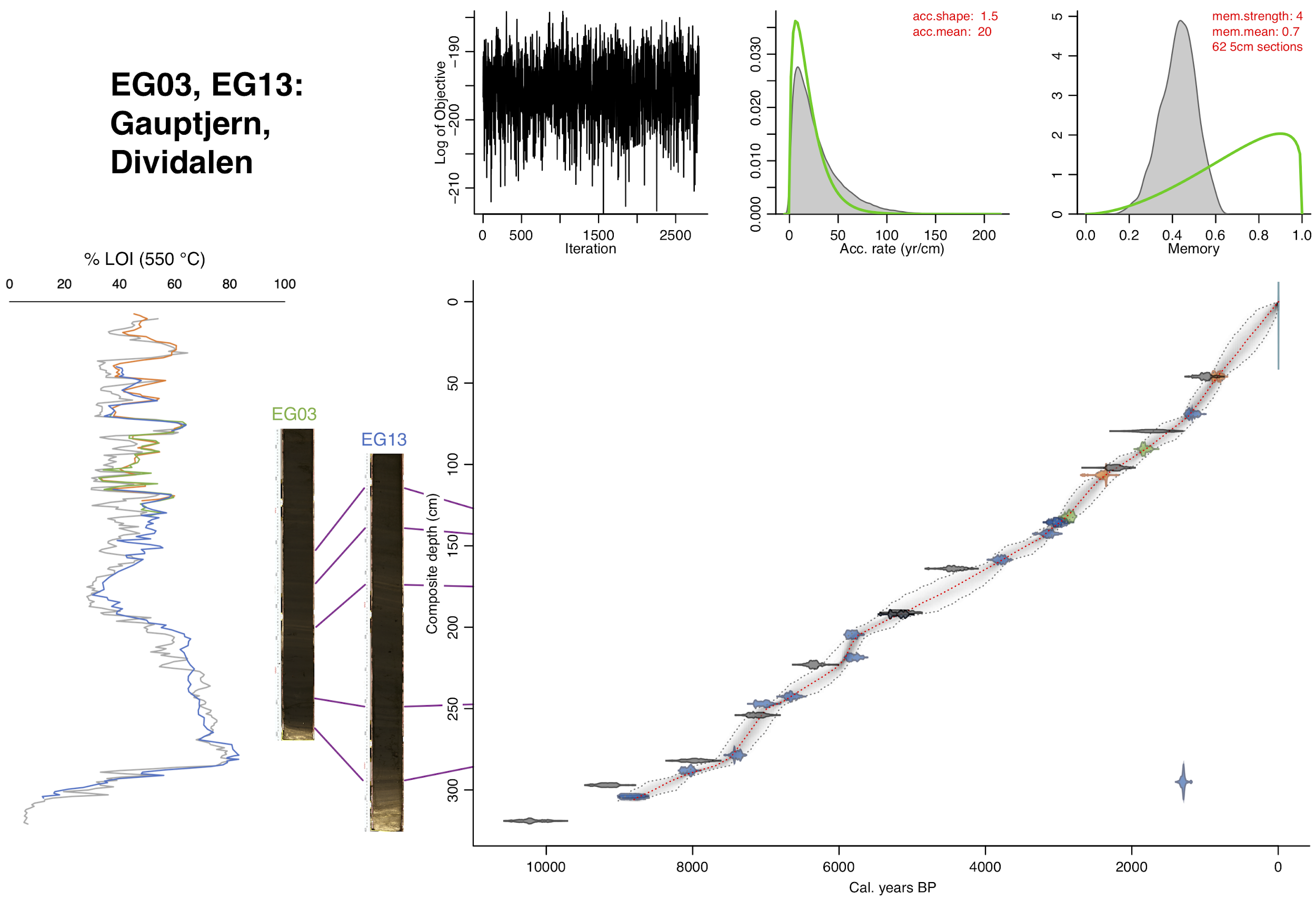

**C**

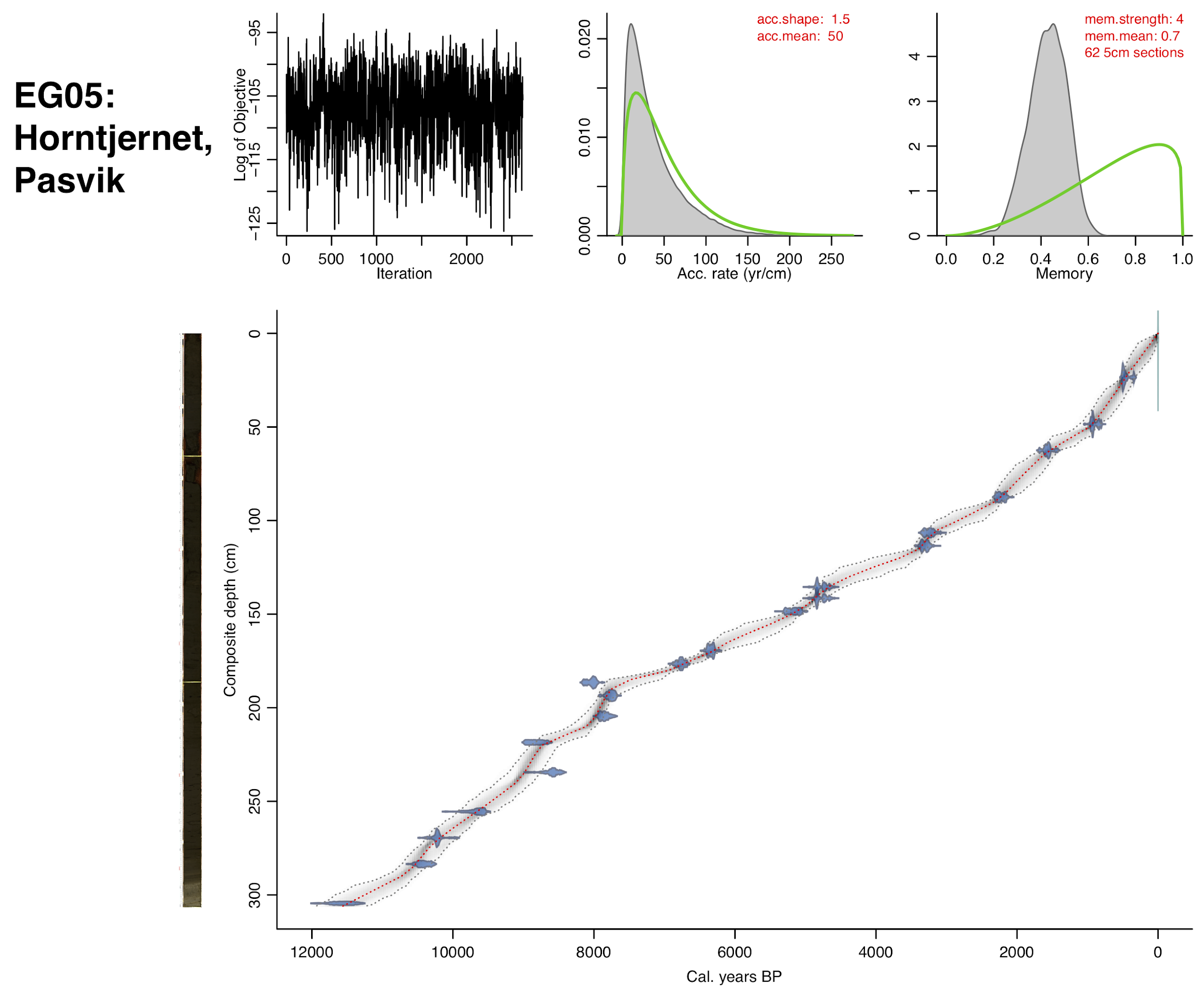

**D**

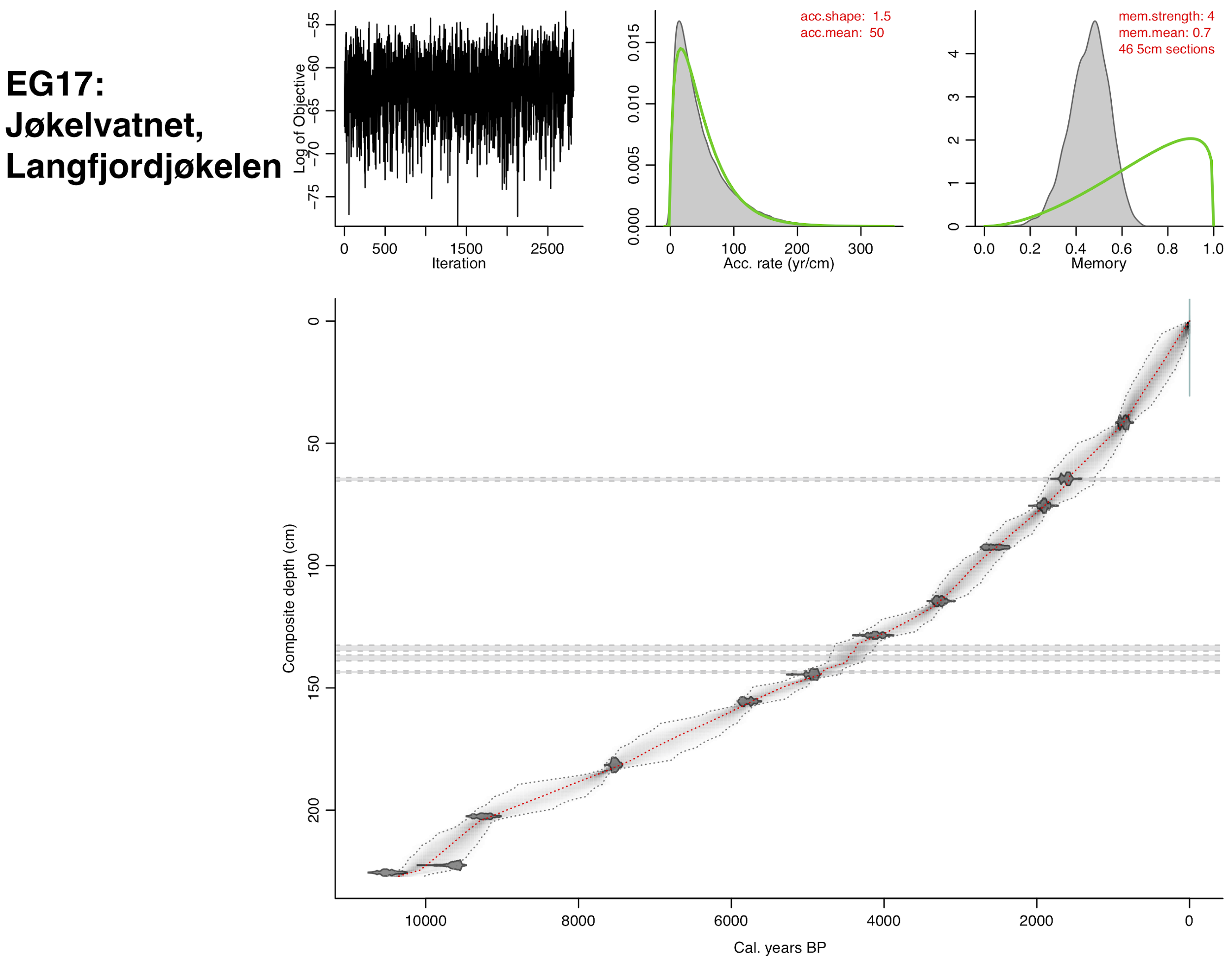

**E**

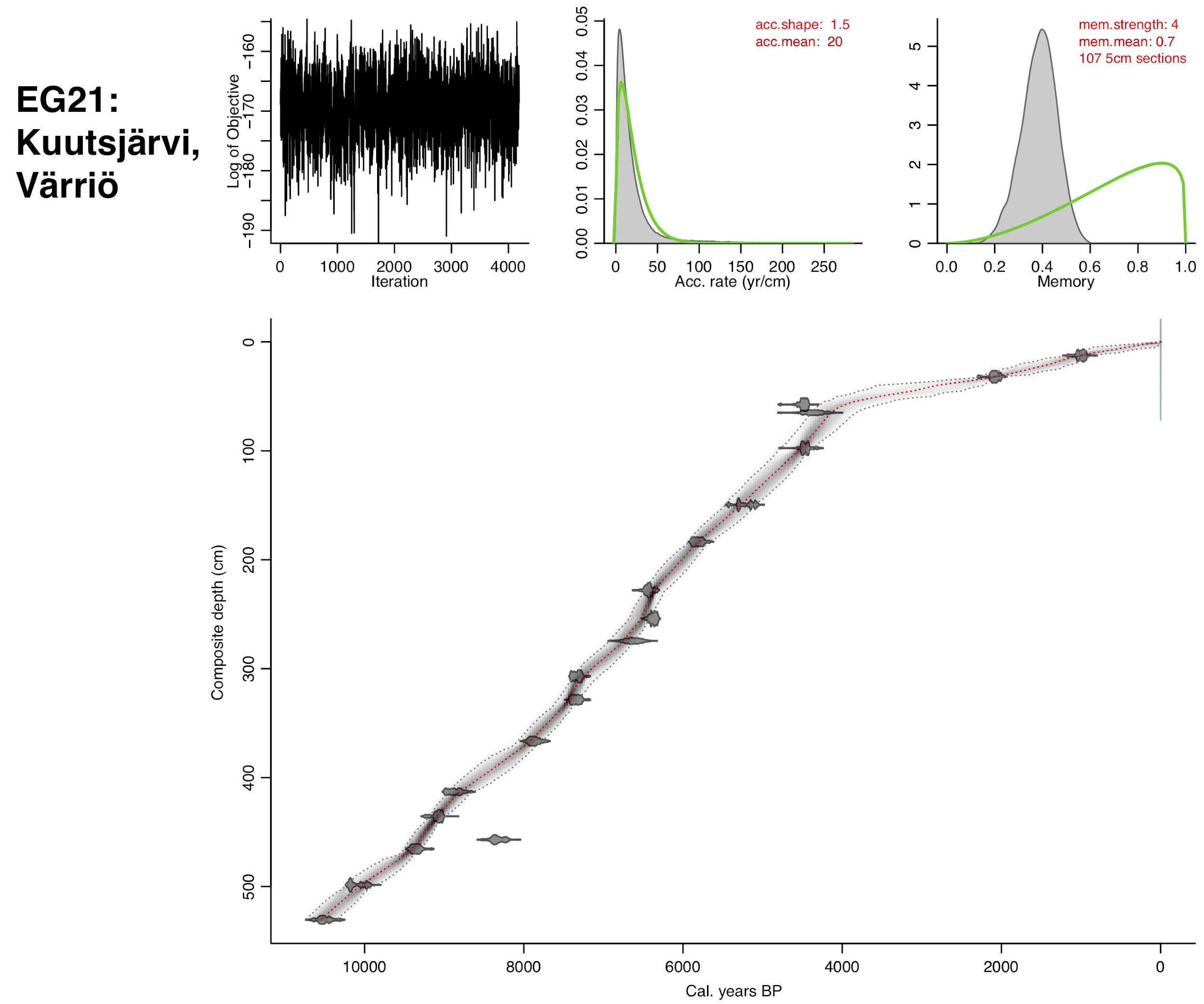

**F**

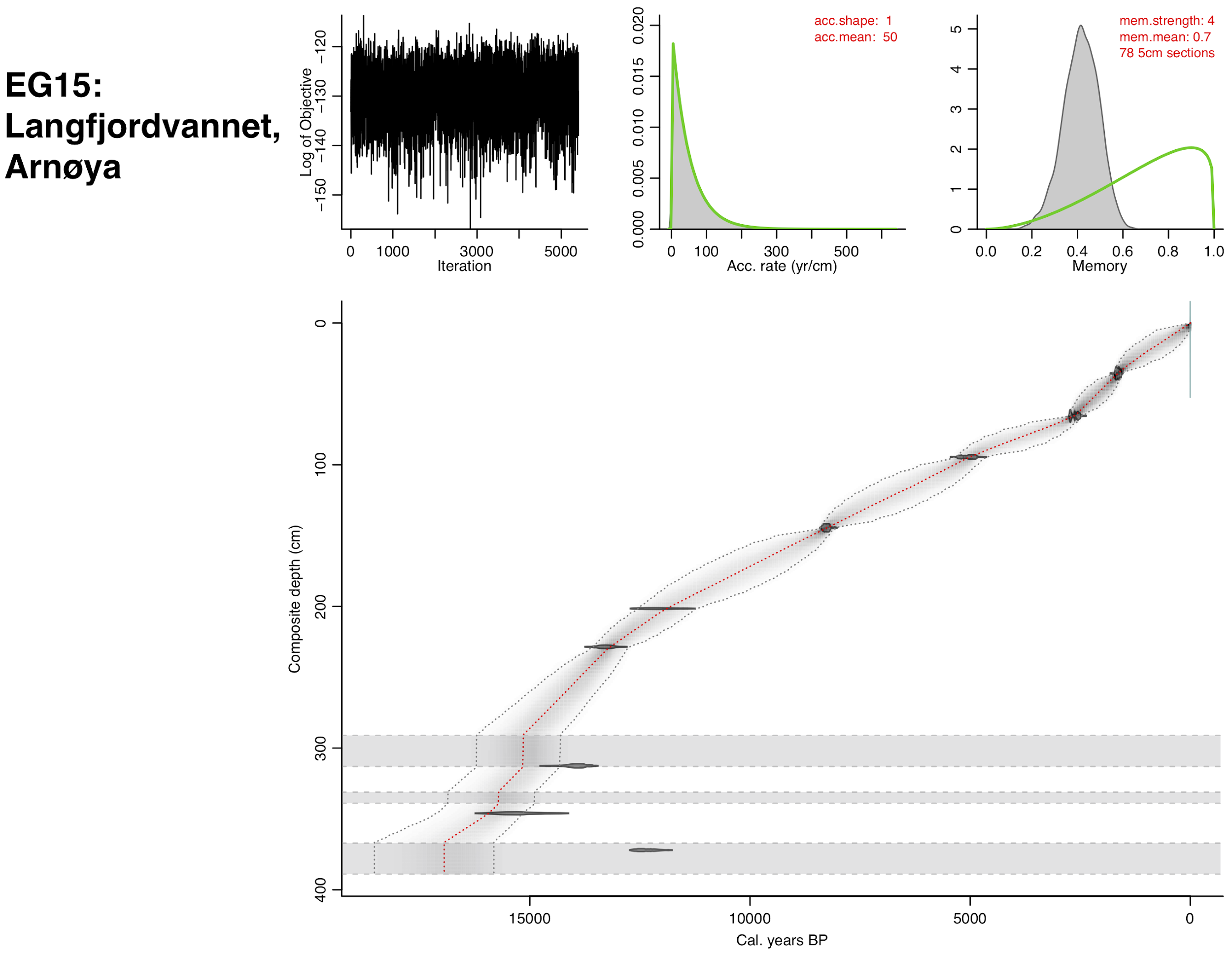

**G**

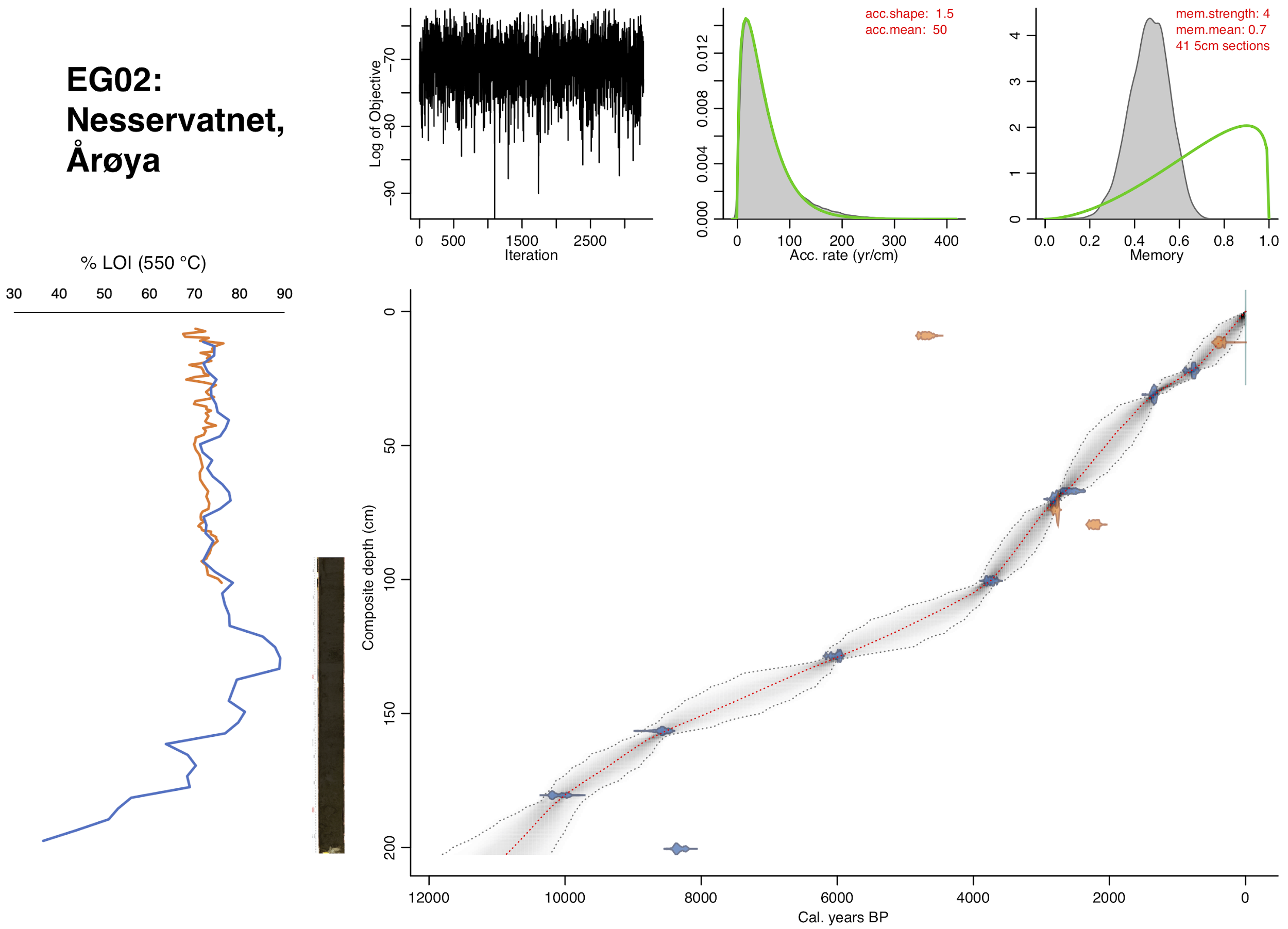

**H**

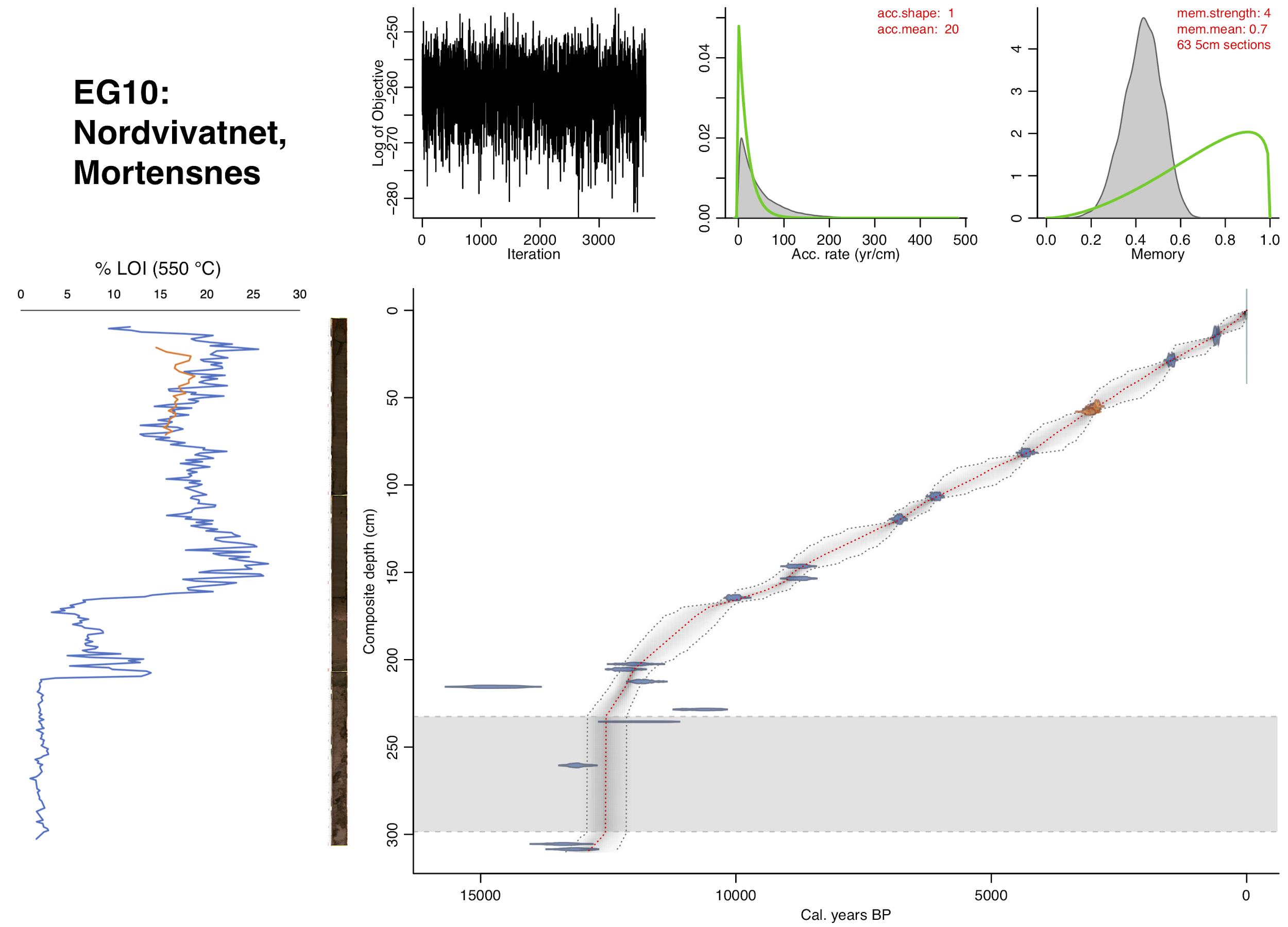

**I**

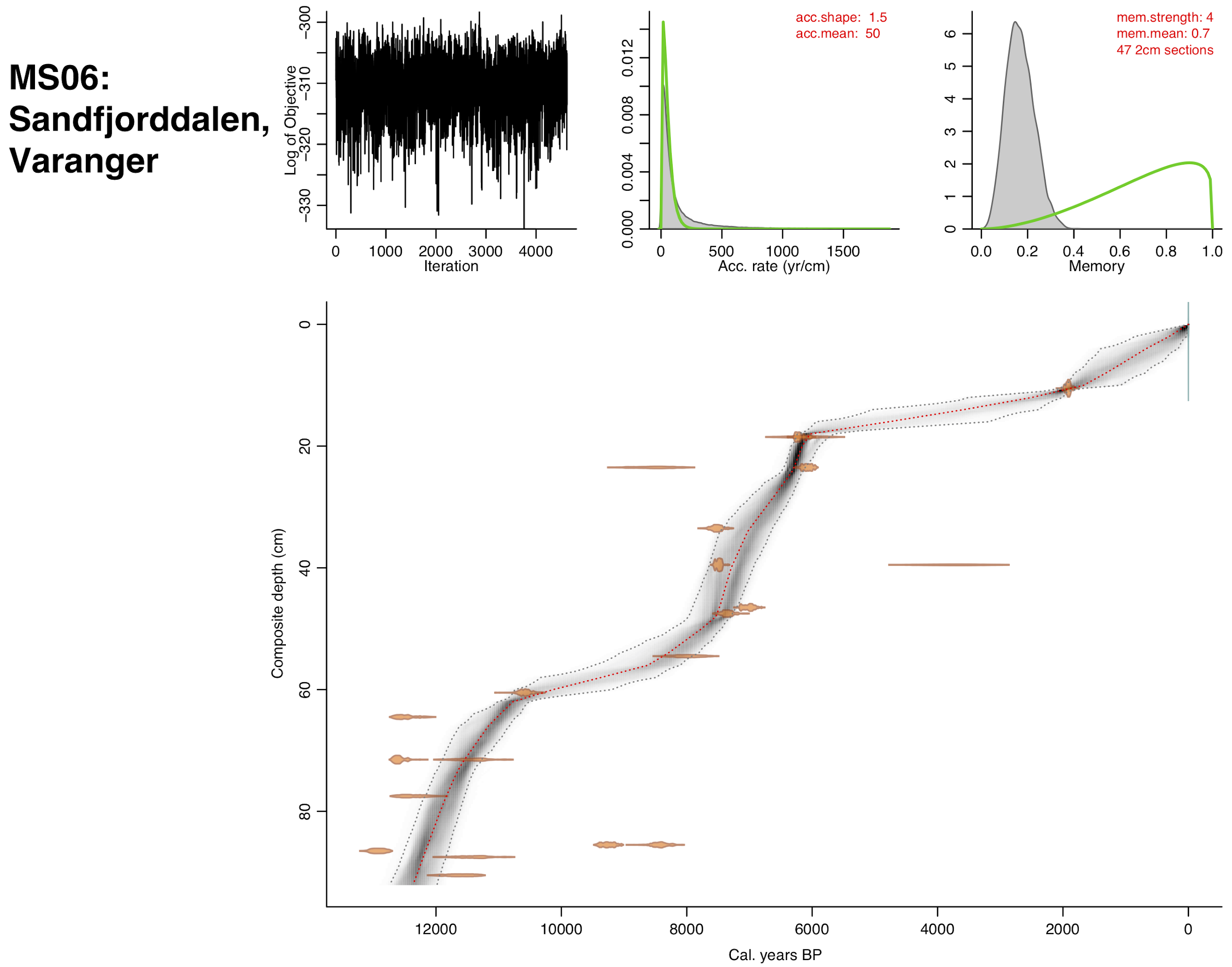

**J**

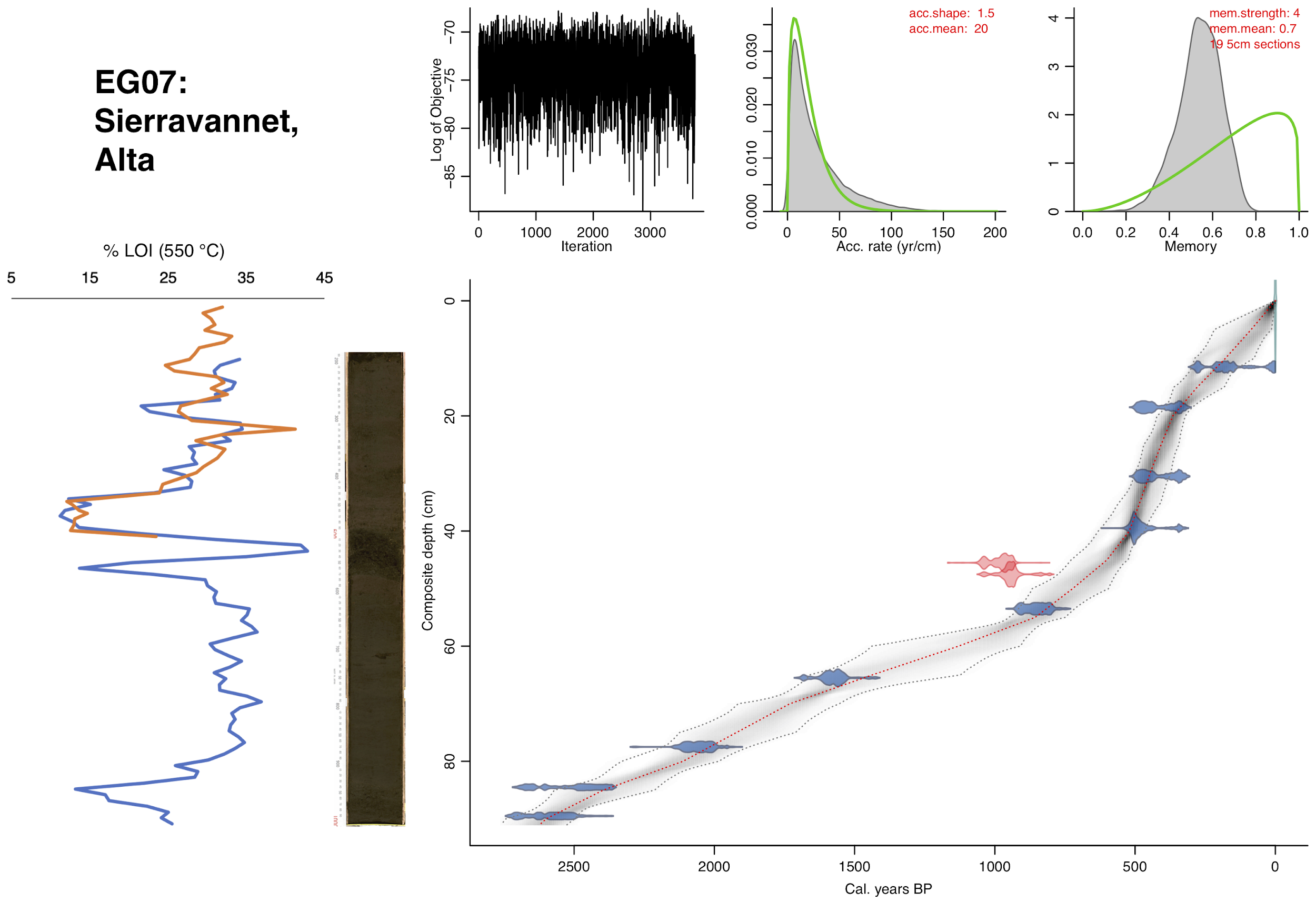

**Fig. S1. Alignments of core LOI, high-res. imagery, and Bayesian age-depth models.**

Colours for calibrated radiocarbon dates and loss-on-ignition (LOI) values are based on Nesje core (dark blue), Multisampler core (orange), or 'other' core (grey). (**A**) EG11: Eaštorjávri South, Ifjord; (**B**) EG03,EG13: Gauptjern, Dividalen (grey dates and LOI data from Jensen & Vorren (*47*)); (**C**) EG05: Horntjernet, Pasvik; (**D**) EG17: Jøkelvatnet, Langfjordjøkelen (grey dates from Wittmeier *et al*. (*55*)); (**E**) EG21: Kuutsjärvi, Värriö (grey dates from Bogren (*56*) or are published in this study); (**F**) EG15: Langfjordvannet, Arnøya (grey dates from Otterå (*31*)); (**G**) EG02: Nesservatnet, Årøya; (**H**) EG10: Nordvivatnet, Mortensnes; (**I**) MS06: Sandfjorddalen, Varanger; (**J**) EG07: Sierravannet, Alta. For (**B**), Nesje cores EG03 and EG13 are in green and dark blue, respectively, and purple lines show five layers anchored by visible stratigraphy. For (**D**), two excluded dates are in red (see Results). For (**E**), (**G**), and (**H**), grey dashed lines and fill in the age-depth model panels illustrate slumps (see Results and Data set S2).

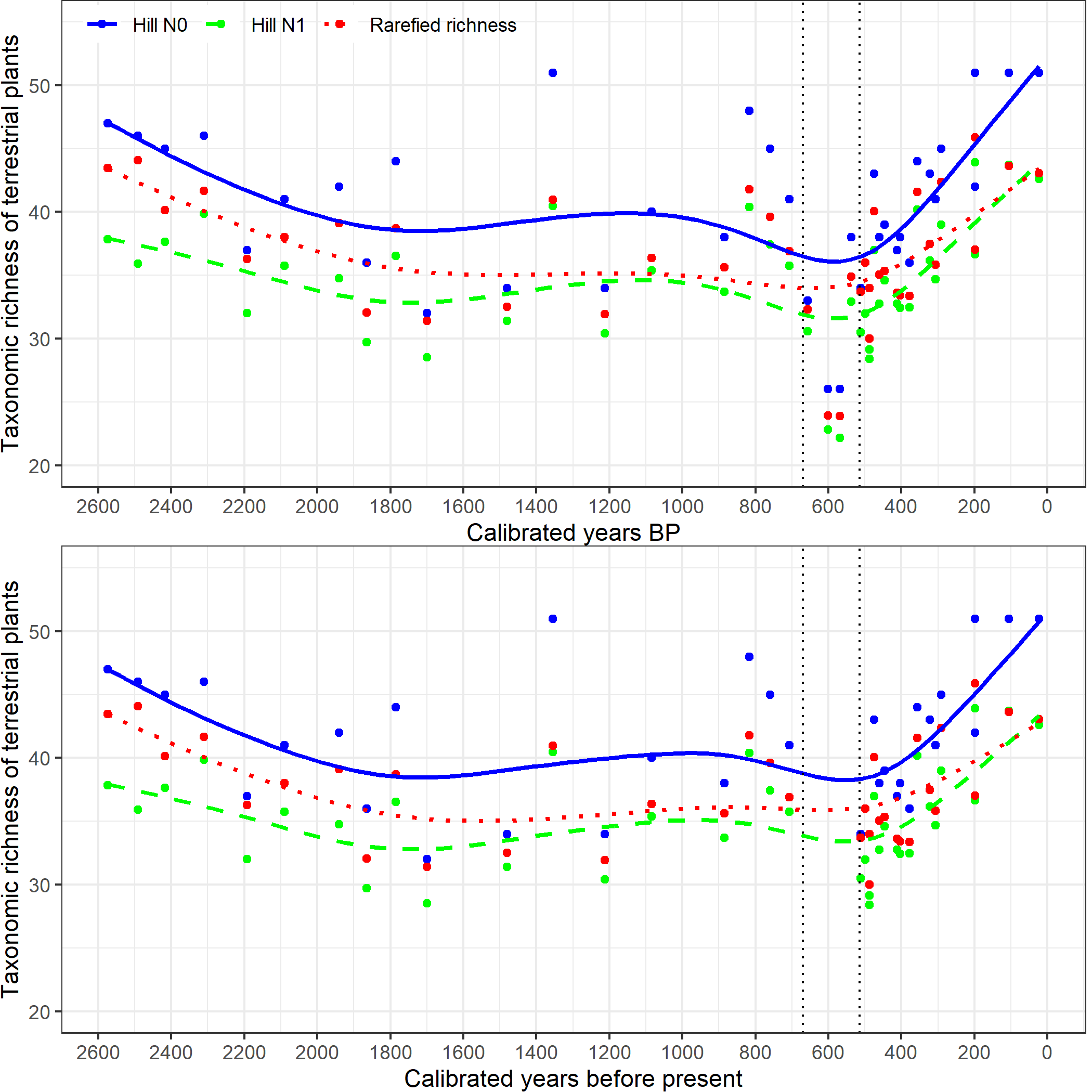

**Fig. S2. A potential flood event does not impact the Sierravannet diversity trend**.

A putative flood event is inferred at ca. 48-40 cm (equivalent to ca. 670-515 years before present) between vertical dotted bars. The richness pattern is unaffected by the removal (lower panel) or inclusion (upper panel) of four samples that fall within this flood event window.

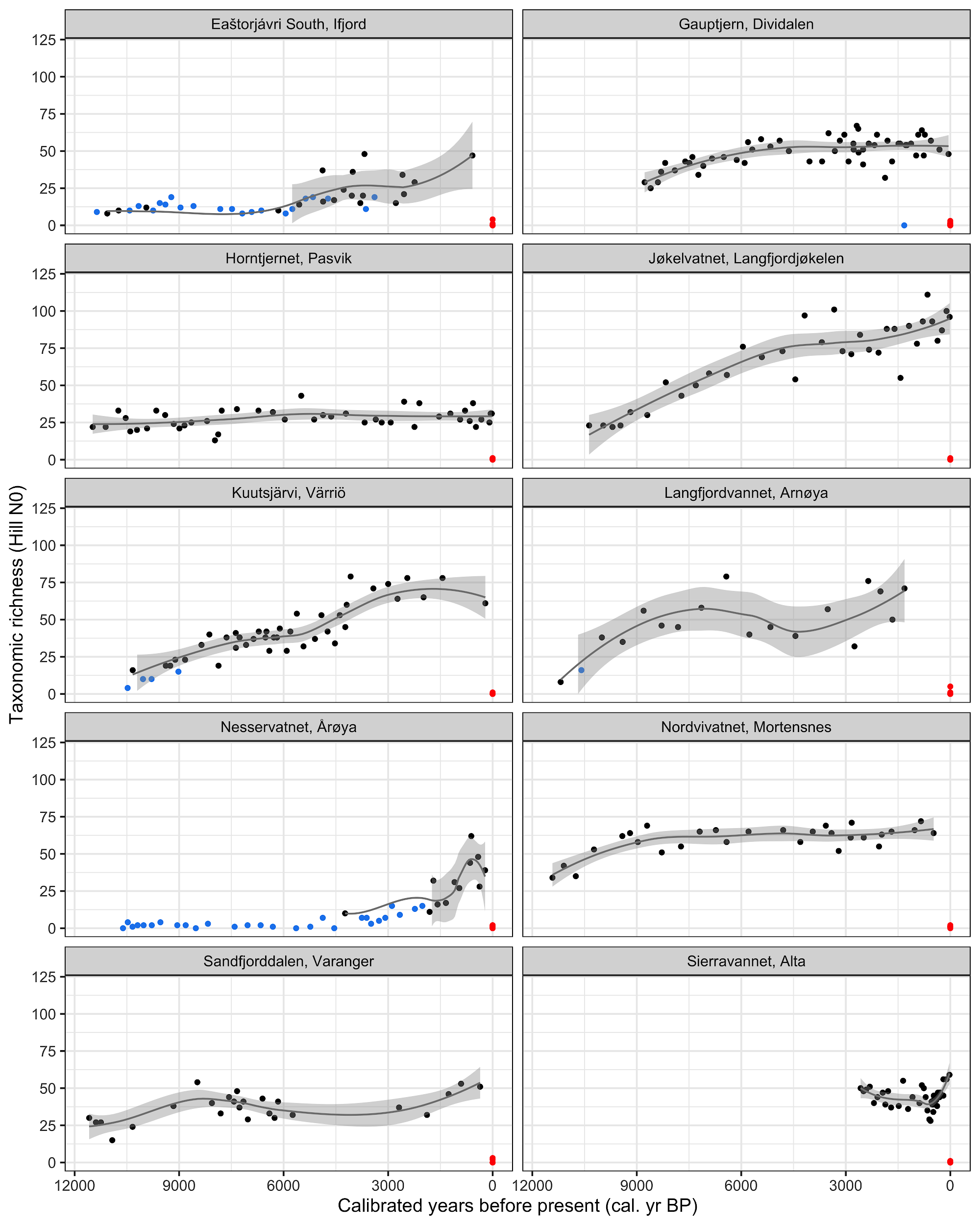

**Fig. S3. Observed taxonomic richness in each sample by lake and time including samples not passing quality controls***.* Samples are coloured: black, passed quality control; blue, failed quality control; whereas red, negative controls. Lines are loess smoothing and shading is one standard error of samples that passed QC.

**A**

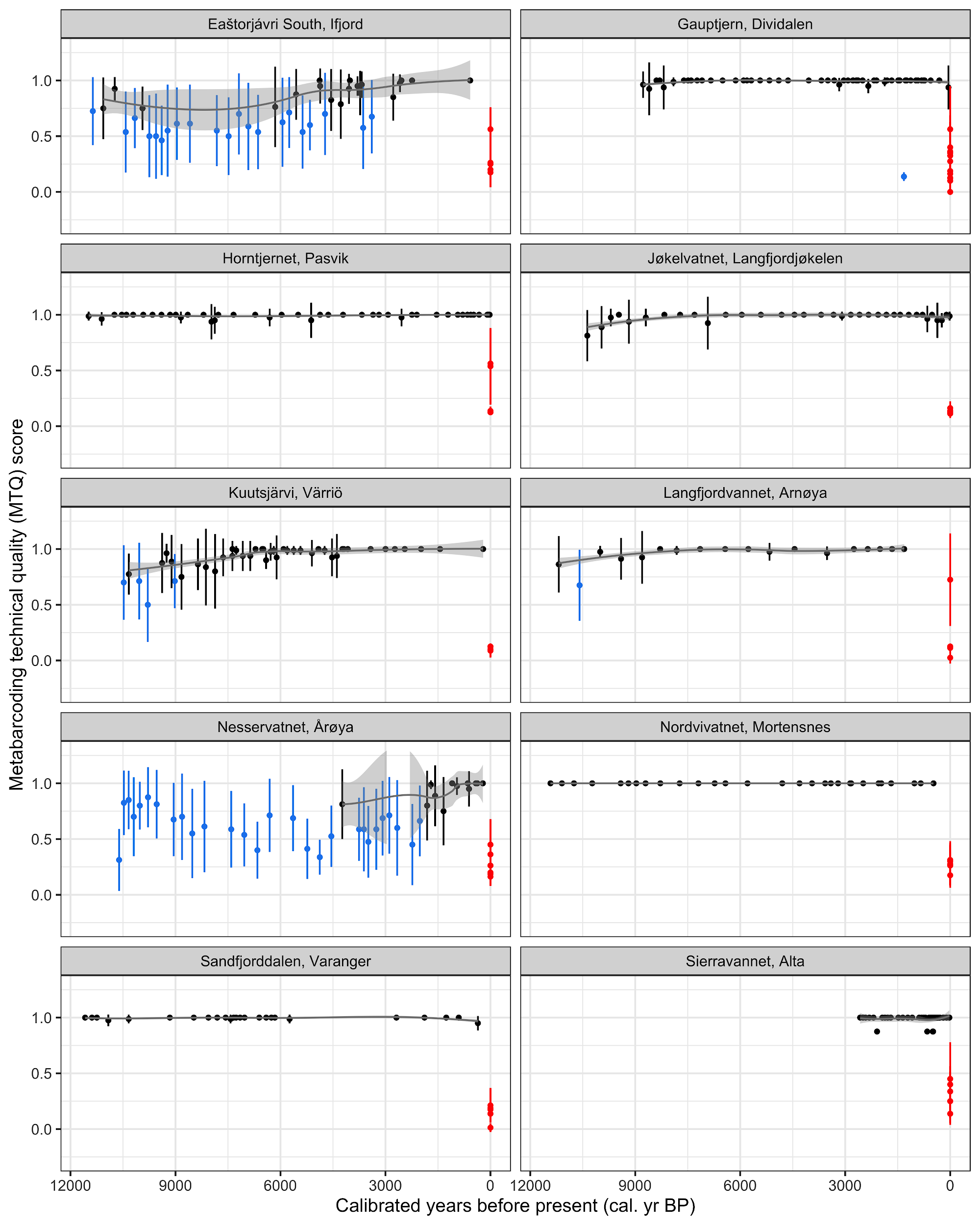

**B**

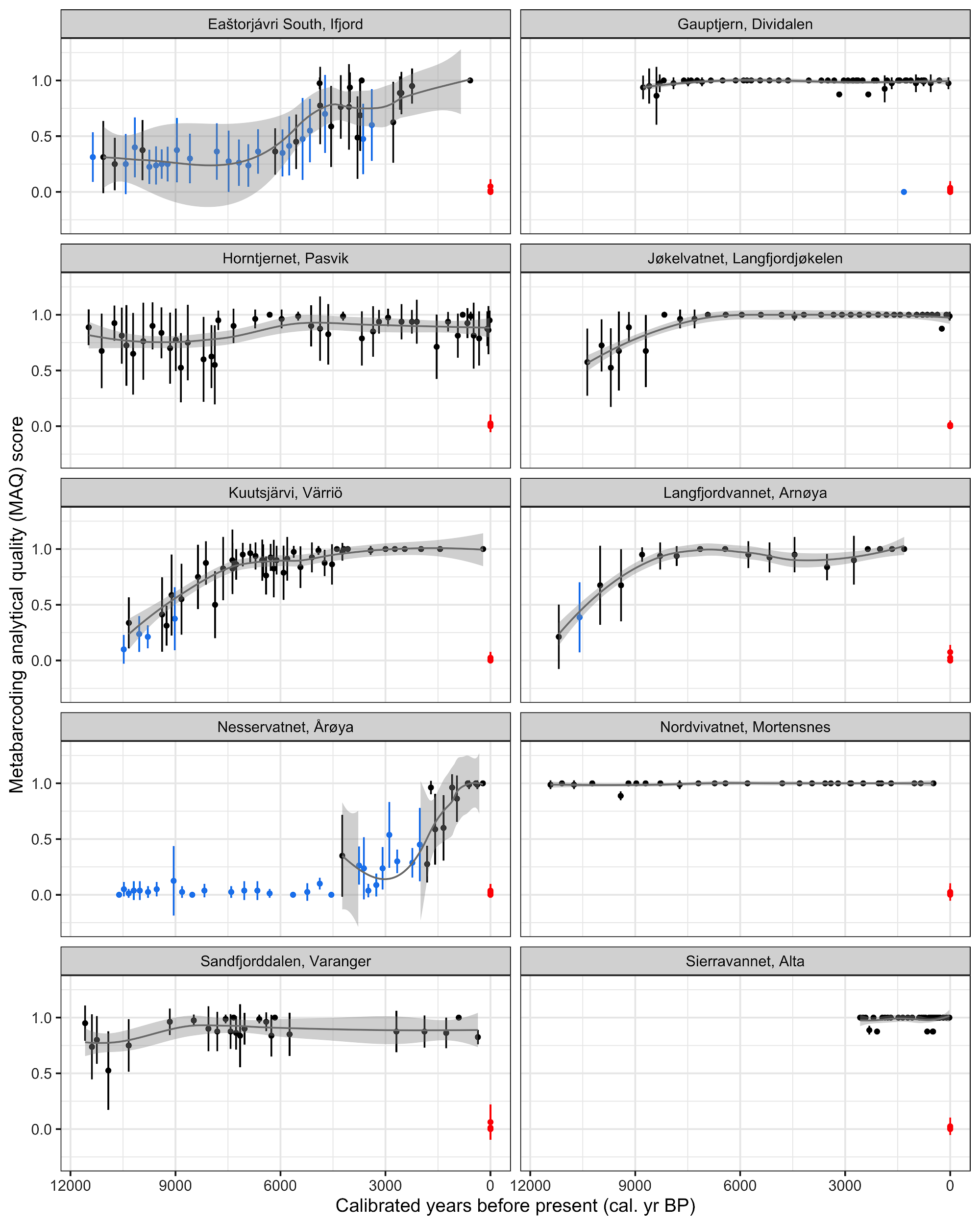

**C**

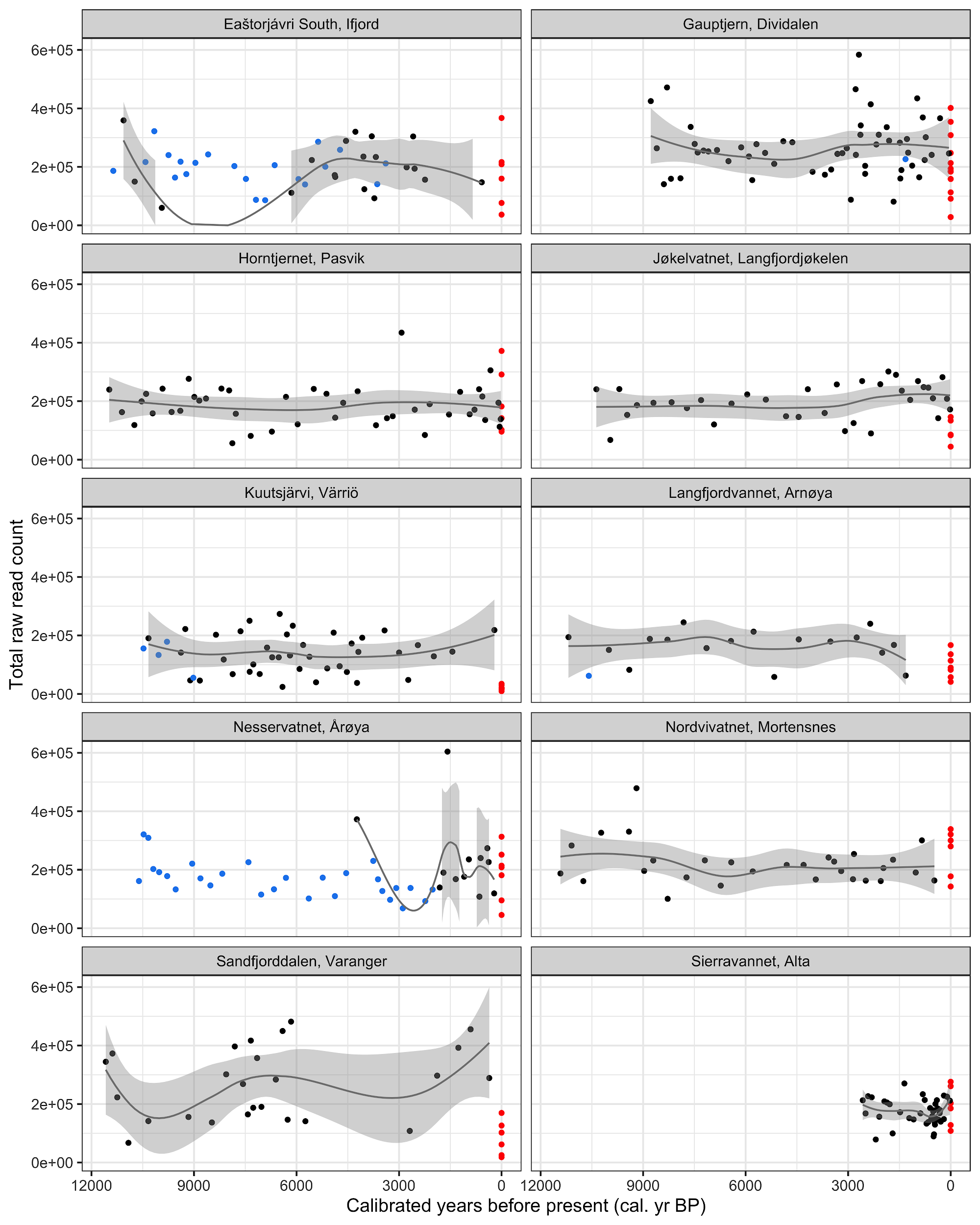

**D**

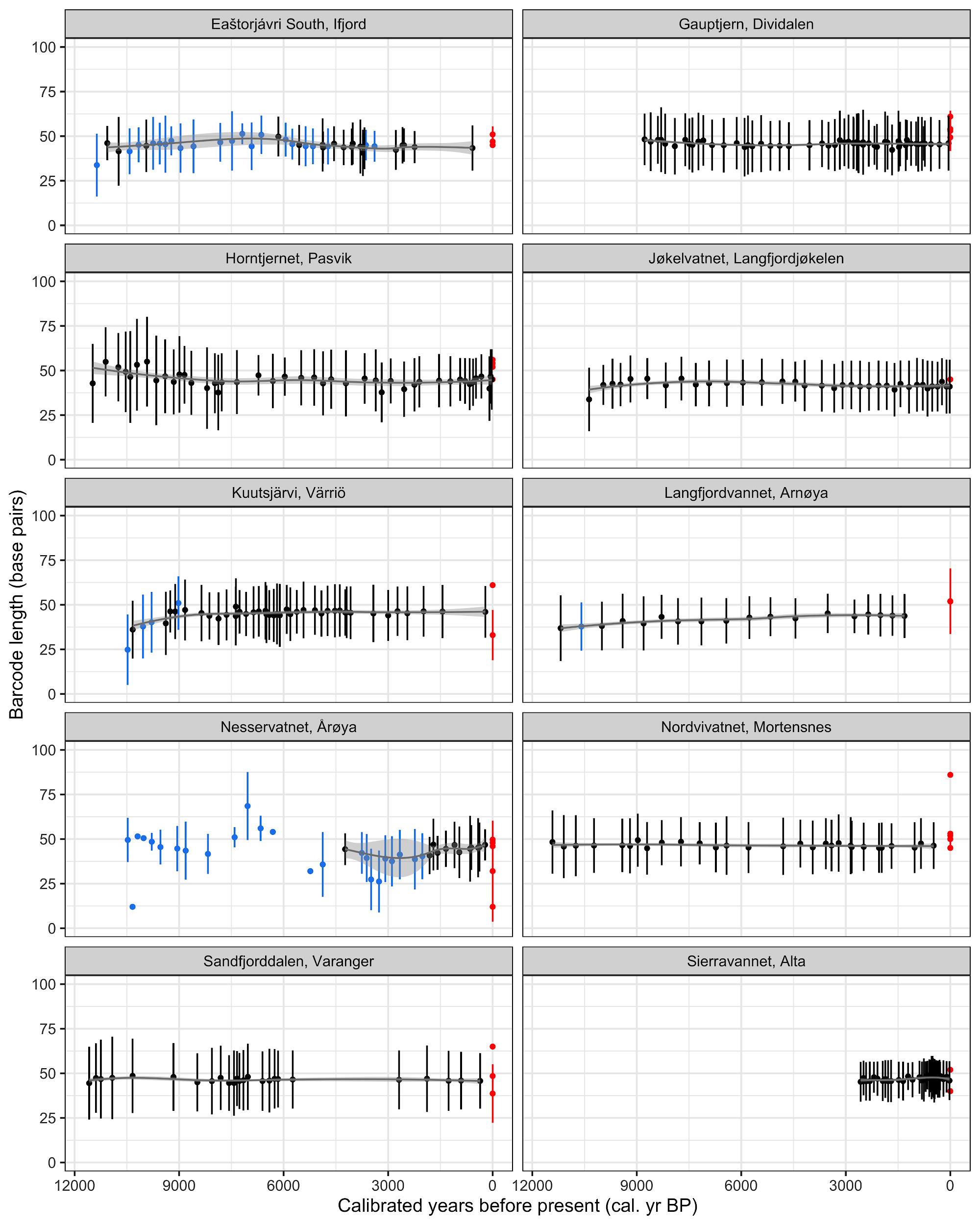

**E**

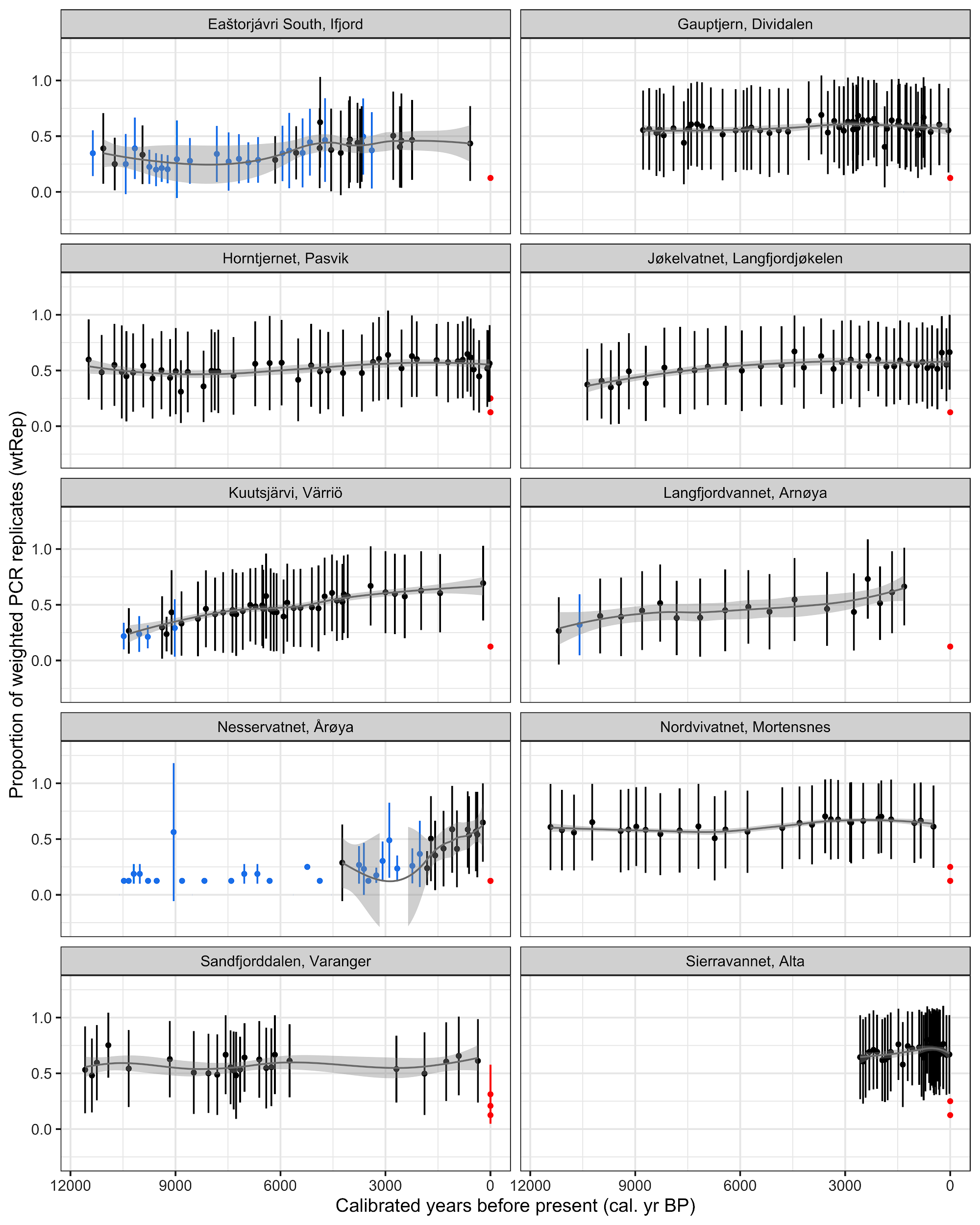

**F**

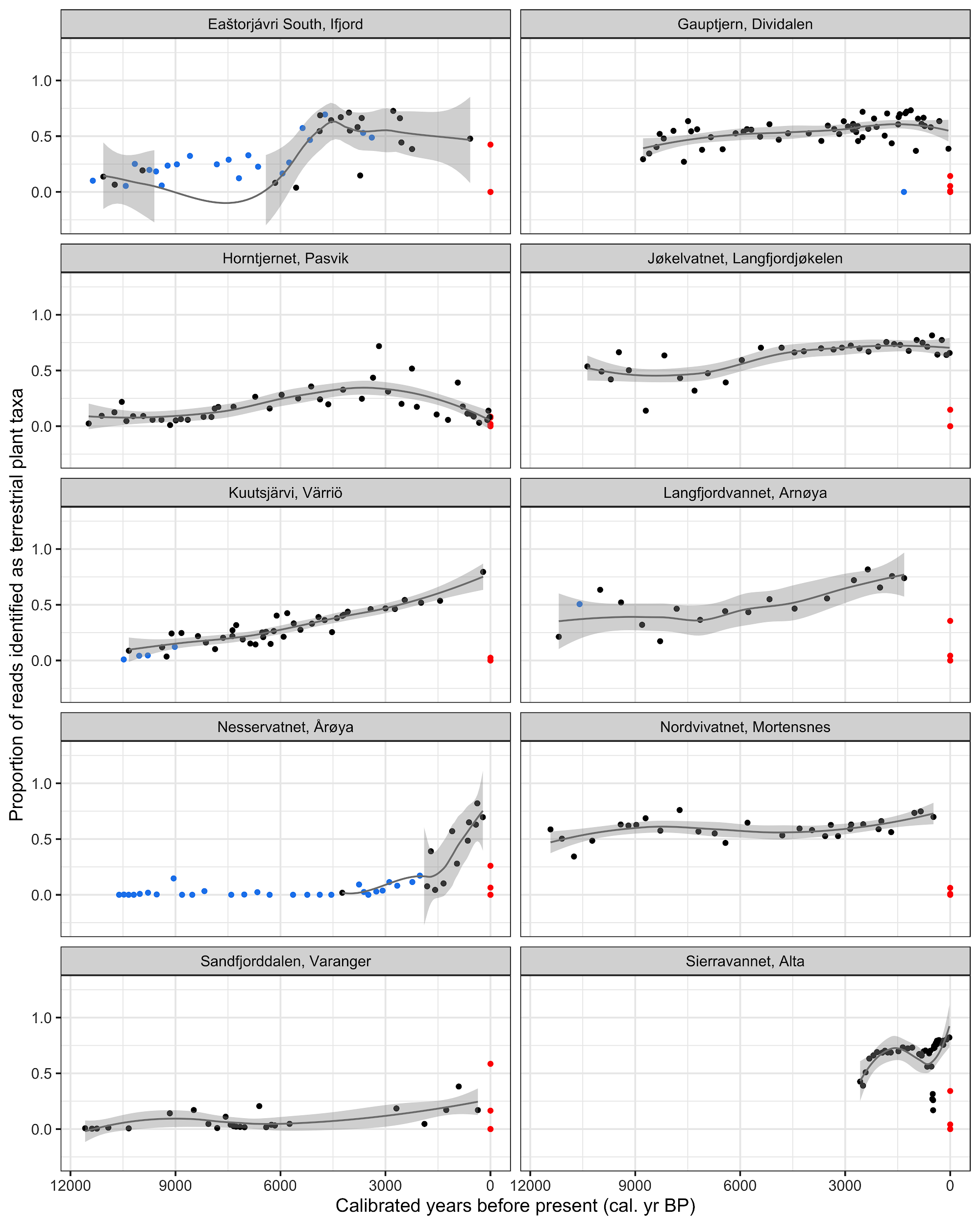

**Fig. S4. Six measures of *sed*aDNA data quality by lake and time**. The six measures are: (**A**) *Metabarcoding technical quality (MTQ)* score, which is an assessment of metabarcoding success and is the proportion of positive PCR detections across the 10 most read-abundant sequences prior to taxonomic identification; (**B**) *metabarcoding analytical quality (MAQ)* score, which is the proportion of positive PCR detections across the 10 most read-abundant and taxonomically-identified sequences on a per sample basis; (**C**) the total raw read count, summed across all eight PCR replicates; (**D**) the proportion of weighted PCR replicates (*wtRep*) averaged across all taxa within each sample; (**E**) the length of retained DNA barcodes, and (**F**) the proportion of raw reads assigned to terrestrial plant taxa. For (**A**), (**B**), (**D**), and (**E**), we show the mean value and one standard deviation across the eight PCR replicates. For (**C**) and (**F**), we show values based on all PCR replicates combined. Samples are coloured: black, passed quality control; blue, failed quality control; whereas red, negative controls. Lines are loess smoothing and shading is one standard error of samples that passed QC.

**A**

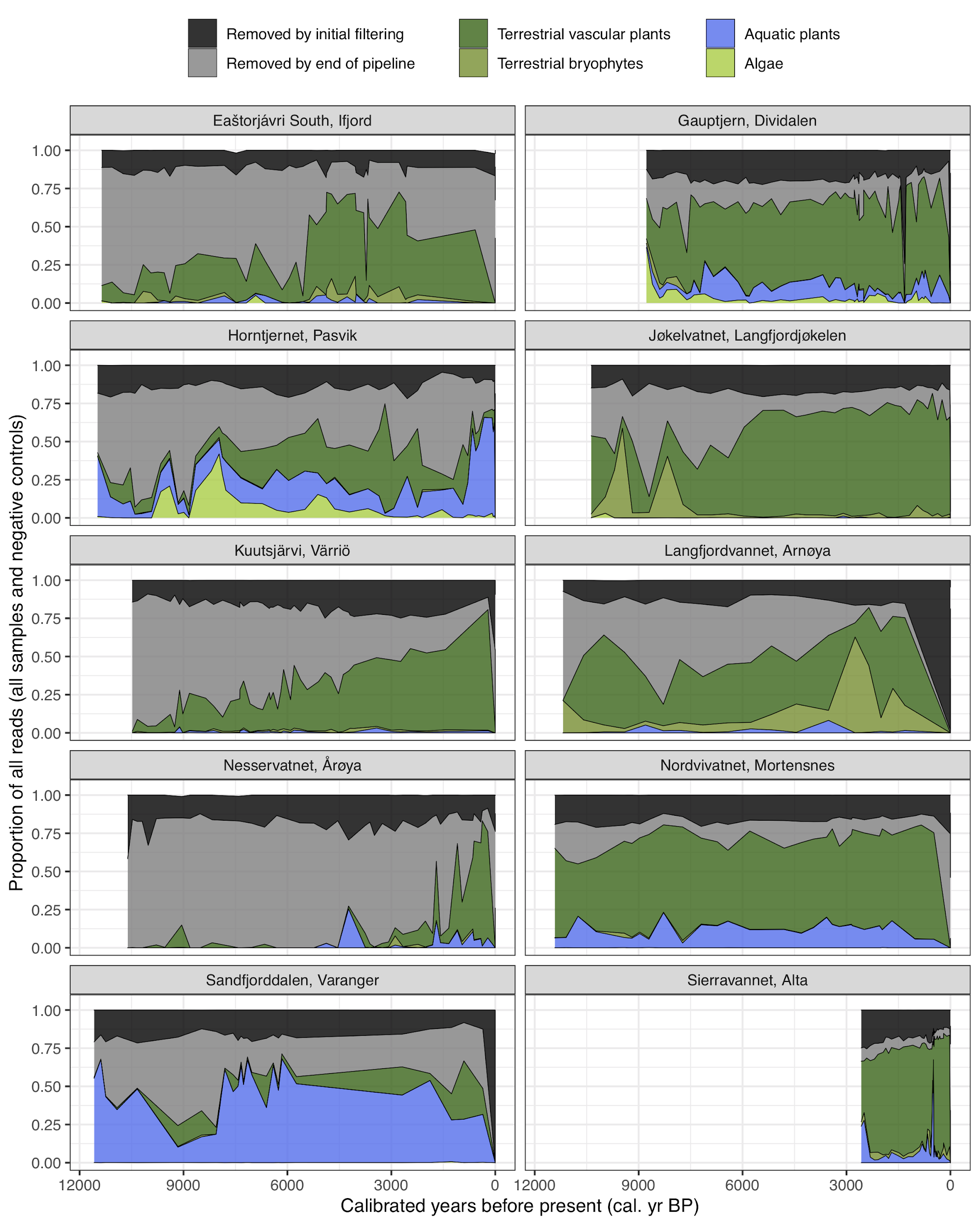

**B**

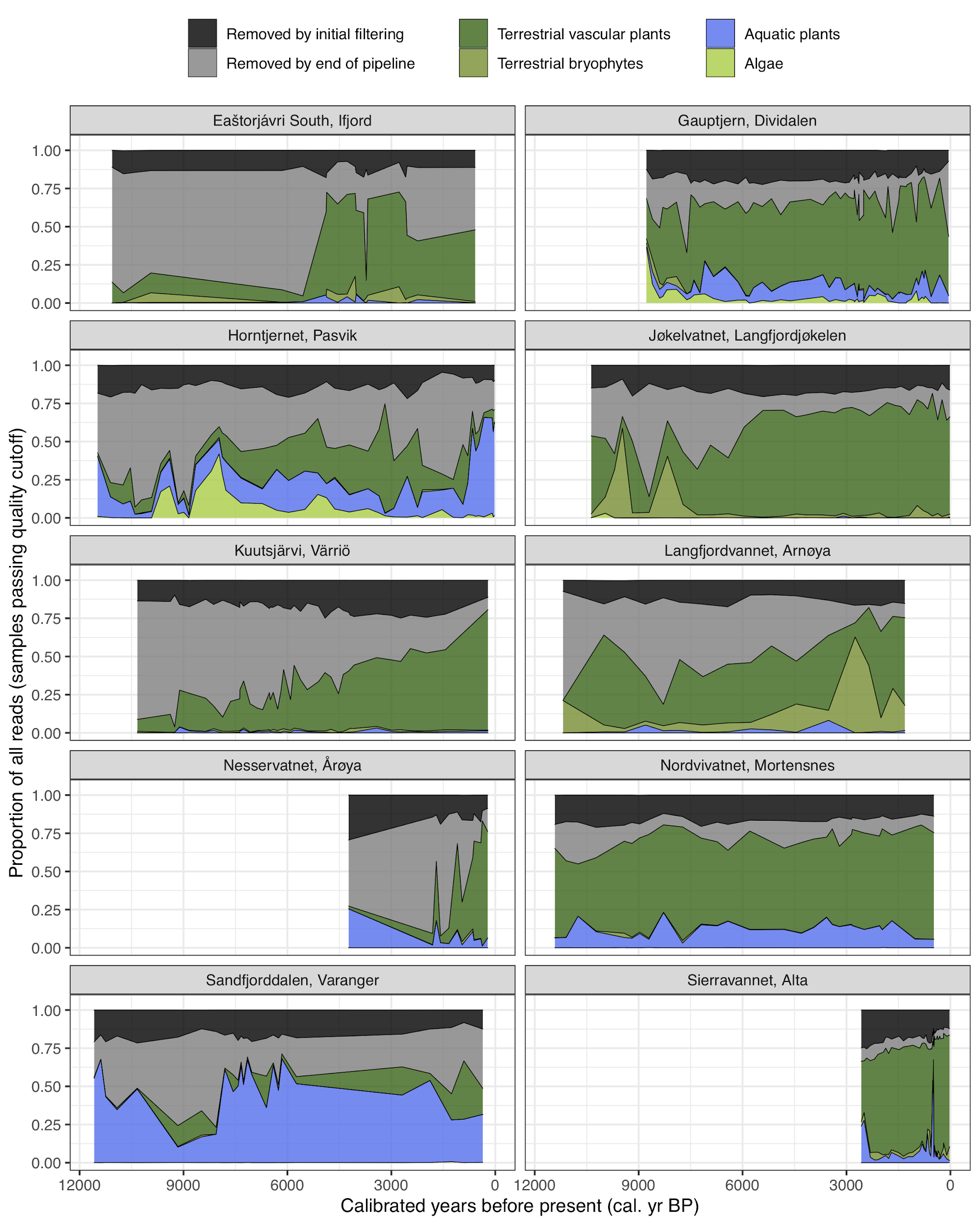

**Fig. S5. The assignments of reads processed by the bioinformatic pipeline**. Reads were summed across all eight PCR replicates and divided into (1) removed by initial filtering (minimum length of 10 bp), (2) removed by the end of the pipeline (unidentified and/or denoised reads), or (3) remaining identified reads that were classified based on broad floral groups. Data are shown for (**A**) all samples and controls prior to quality control, and (**B**) only samples that passed quality control.

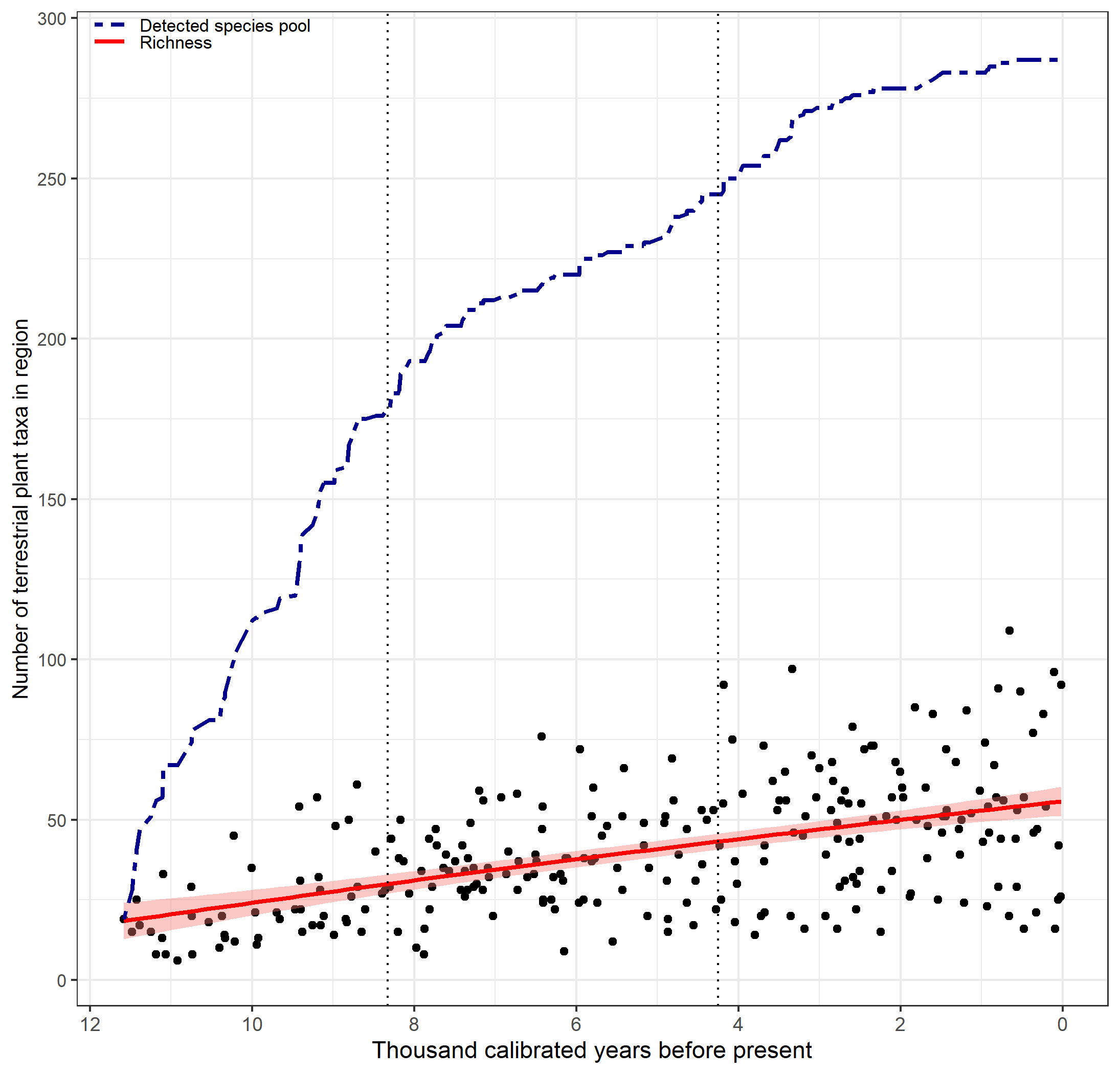

**Fig. S6. The accumulated regional species pool and taxonomic richness of each sample across the Holocene of northern Fennoscandia excluding two temporally-short records from Nesservatnet (EG02) and Sierravannet (EG07)**. The accumulation of the detected regional species pool (defined as cumulative number of taxa; double-dashed line) as well as number of taxa detected per sample (n=264) along with the 95% confidence interval (pink shading) of the fitted line (solid red line) based on a generalized additive model.

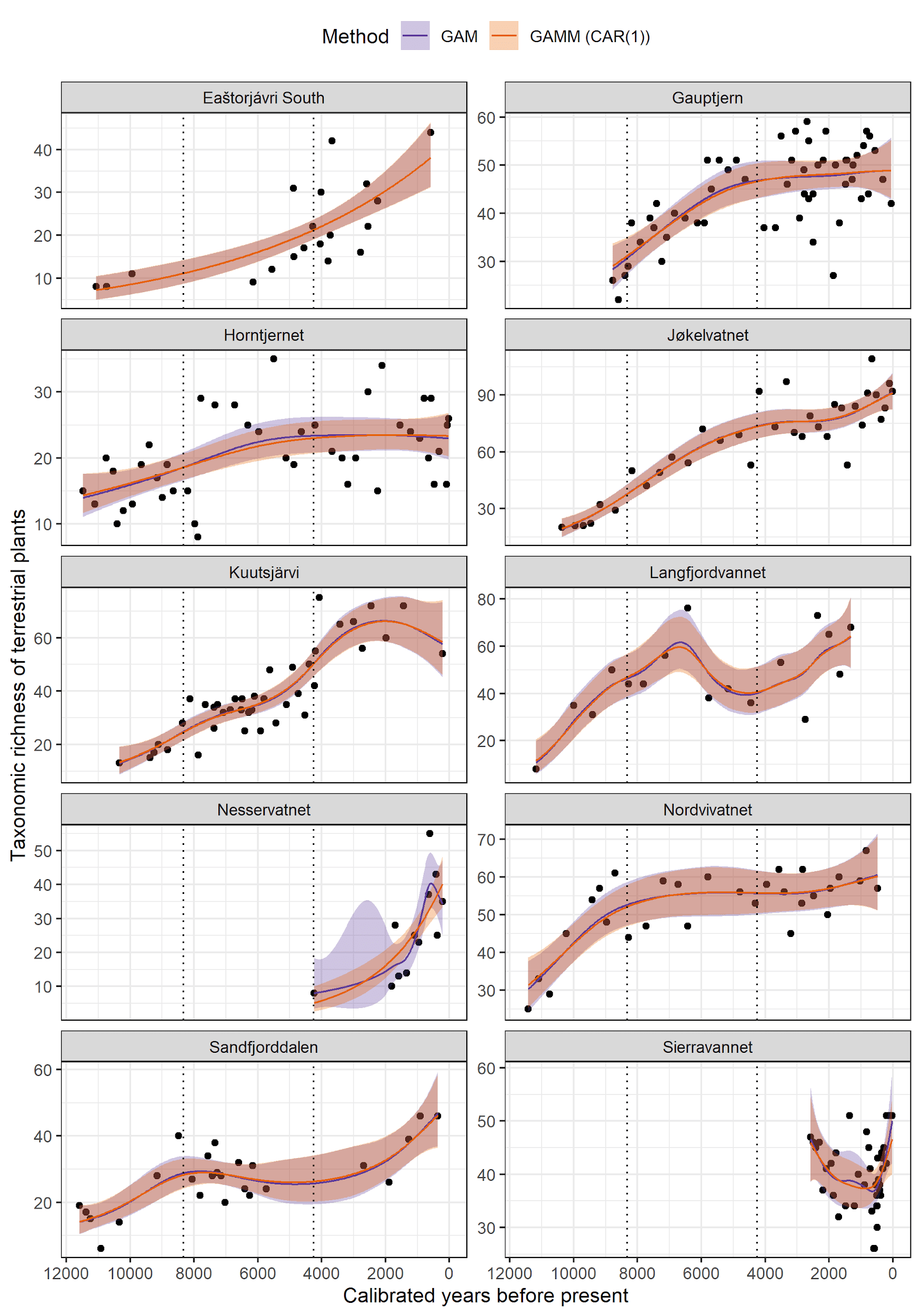

**Fig. S7. Comparison between GAM and GAMM(CAR(1)) models of taxonomic richness through time**. The fitted lines and their 95% confidence intervals are based on generalized additive model (GAM), and generalized additive mixed model (GAMM) with a continuous time first-order autoregressive process (CAR(1)). The Early (12.0-8.3 ka), Middle (8.3-4.25 ka), and Late Holocene (4.25-0.0 ka) periods are separated by vertical dotted lines.

Table S1. Summary of all data used or generated in this study. *Totals include previously published data. **Excluding duplicates and negative controls. Full data are given in Data sets S3-S4.

|  |  |  |  |  |  | **Taxonomic richness** | | | | |
| --- | --- | --- | --- | --- | --- | --- | --- | --- | --- | --- |
|  |  |  |  | **DNA samples**** | | **Observed (Hill-N0)** | | | **Hill-N1** | |
| **Code** | **Locality** | **Radiocarbon dates** | **LOI samples** | **All** | **Passing QC** | **Total** | **Mean** | **Stdev** | **Mean** | **Stdev** |
| EG02 | Nesservatnet, Årøya | 13 | 136 | 40 | 12 | 93 | 26.33 | 14.27 | 20.43 | 12.02 |
| EG03, EG13 | Gauptjern, Dividalen | 30* | 558* | 56 | 55 | 163 | 44.09 | 9.03 | 34.82 | 7.39 |
| EG05 | Horntjernet, Pasvik | 20 |  | 44 | 44 | 114 | 20.61 | 6.43 | 15.89 | 5.30 |
| EG07 | Sierravannet, Alta | 11 | 119 | 40 | 40 | 108 | 40.08 | 6.48 | 34.52 | 4.92 |
| EG10 | Nordvivatnet, Mortensnes | 20 | 302 | 29 | 29 | 160 | 52.31 | 9.95 | 42.89 | 8.64 |
| EG11 | Eaštorjávri South, Ifjord | 15 | 95 | 40 | 19 | 89 | 21.00 | 10.86 | 14.87 | 7.78 |
| EG15 | Langfjordvannet, Arnøya | 9* |  | 18 | 17 | 154 | 46.82 | 17.50 | 35.43 | 14.83 |
| EG17 | Jøkelvatnet, Langfjordjøkelen | 12* |  | 35 | 35 | 200 | 65.54 | 24.47 | 52.40 | 20.49 |
| EG21 | Kuutsjärvi, Värriö | 19 |  | 44 | 40 | 138 | 38.83 | 16.51 | 30.05 | 14.06 |
| MS06 | Sandfjorddalen, Varanger | 21 |  | 25 | 25 | 97 | 27.44 | 9.79 | 21.77 | 8.06 |
|  | **Totals** | **170*** | **1210*** | **371** | **316** |  |  |  |  |  |

Table S2. Correlations between observed and rarefied taxonomic richness for each lake. R, Pearson product moment correlation; 95% confidence interval is for R.

| **Code** | **Lake** | **t** | **df** | **p-value** | **R** | **R.95.low** | **R.95.high** |
| --- | --- | --- | --- | --- | --- | --- | --- |
| EG02 | Nesservatnet | 12.04 | 10 | 0 | 0.97 | 0.88 | 0.99 |
| EG03/13 | Gauptjern | 37.55 | 53 | 0 | 0.98 | 0.97 | 0.99 |
| EG05 | Horntjernet | 15.04 | 42 | 0 | 0.92 | 0.85 | 0.95 |
| EG07 | Sierravannet | 19.98 | 38 | 0 | 0.96 | 0.92 | 0.98 |
| EG10 | Nordvivatnet | 50.09 | 27 | 0 | 0.99 | 0.99 | 1.00 |
| EG11 | Eaštorjávri South | 37.80 | 17 | 0 | 0.99 | 0.98 | 1.00 |
| EG15 | Langfjordvannet | 33.79 | 15 | 0 | 0.99 | 0.98 | 1.00 |
| EG17 | Jøkelvatnet | 36.43 | 33 | 0 | 0.99 | 0.98 | 0.99 |
| EG21 | Kuutsjärvi | 26.52 | 38 | 0 | 0.97 | 0.95 | 0.99 |
| MS06 | Sandfjorddalen | 6.77 | 23 | 0 | 0.82 | 0.62 | 0.92 |

Table S3. Summary of generalized additive models (GAMs), and generalized additive mixed models (GAMMs) with a continuous time first-order autoregressive (CAR(1)) process. Median calibrated age of samples was treated as the predictor and taxonomic richness as the response variables. edf: effective degrees of freedom; Dev.exp: deviance explained; Phi (ɸ): autocorrelation coefficient; adj.R.sq.: adjusted R square.

| **Code** | **Lake** | **Model** | **edf** | **Tests** | **Test score** | **p-value** | **Phi (ɸ)** | **Dev.exp (%)** | **adj.R.sq** |
| --- | --- | --- | --- | --- | --- | --- | --- | --- | --- |
| EG02 | Nesservatnet | GAM | 3.92 | ꭕ2 | 51.34 | 0 | NA | 73.48 | 0.56 |
| EG03/13 | Gauptjern |  | 2.98 | ꭕ2 | 47.11 | 0 | NA | 50.9 | 0.46 |
| EG05 | Horntjernet |  | 2.49 | ꭕ2 | 23.03 | 0 | NA | 29.84 | 0.25 |
| EG07 | Sierravannet |  | 3.74 | ꭕ2 | 13.29 | 0.02 | NA | 37.5 | 0.33 |
| EG10 | Nordvivatnet |  | 3.82 | ꭕ2 | 34.62 | 0 | NA | 70.33 | 0.63 |
| EG11 | Eaštorjávri South |  | 1 | ꭕ2 | 46.79 | 0 | NA | 57.36 | 0.51 |
| EG15 | Langfjordvannet |  | 6.76 | ꭕ2 | 58.8 | 0 | NA | 72.9 | 0.45 |
| EG17 | Jøkelvatnet |  | 4.15 | ꭕ2 | 236.27 | 0 | NA | 87.03 | 0.81 |
| EG21 | Kuutsjärvi |  | 4.91 | ꭕ2 | 204.09 | 0 | NA | 81.54 | 0.78 |
| MS06 | Sandfjorddalen |  | 4.09 | ꭕ2 | 52.77 | 0 | NA | 67.39 | 0.64 |
| EG02 | Nesservatnet | GAM (CAR(1)) | 1 | F | 33.87 | 0 | 0.2 | NA | 0.49 |
| EG03/13 | Gauptjern |  | 2.58 | F | 16.83 | 0 | 0.37 | NA | 0.46 |
| EG05 | Horntjernet |  | 1.98 | F | 10.08 | 0 | 0.2 | NA | 0.24 |
| EG07 | Sierravannet |  | 2.73 | F | 3.1 | 0.09 | 0.44 | NA | 0.25 |
| EG10 | Nordvivatnet |  | 3.39 | F | 9.66 | 0 | 0.2 | NA | 0.62 |
| EG11 | Eaštorjávri South |  | 1 | F | 44.4 | 0 | 0.2 | NA | 0.51 |
| EG15 | Langfjordvannet |  | 5.93 | F | 8.91 | 0 | 0.2 | NA | 0.45 |
| EG17 | Jøkelvatnet |  | 3.88 | F | 59.02 | 0 | 0.2 | NA | 0.81 |
| EG21 | Kuutsjärvi |  | 4.53 | F | 43.9 | 0 | 0 | NA | 0.78 |
| MS06 | Sandfjorddalen |  | 3.7 | F | 13.52 | 0 | 0.2 | NA | 0.64 |

Additional data files

Data sets S1 to S7
